# Integrated Metabolomics, Transcriptomics, and Signal Protein Profile Define the Effector Function and Associated Metabotype of Five Polarized Macrophage Phenotypes

**DOI:** 10.1101/2020.03.10.985788

**Authors:** Catherine B. Anders, Tyler M.W. Lawton, Hannah L. Smith, Jamie Garret, Margaret M. Doucette, Mary Cloud B. Ammons

**Author notes:** Corresponding Author: Mary Cloud B. Ammons Ph.D.

## Abstract

Macrophages (MΦs) display remarkable plasticity and the ability to activate diverse responses to a host of intracellular and external stimuli. Despite extensive characterization of M1 MΦs and a broad set of M2 MΦs, comprehensive characterization of metabolic shifts driving MΦ activation remains. Herein, we utilized an *ex vivo* model to produce six MΦ functional phenotypes. Isolated CD14+ PBMCs were differentiated into resting M0 MΦs, and then polarized into M1 (IFN-γ/LPS), M2a (IL-4/IL-13), M2b (IC/LPS), M2c (IL-10), and M2d (IL-6/LIF) MΦs. The MΦs were profiled using a bioanalyte matrix of four cell surface markers, ∼50 secreted proteins, ∼800 expressed myeloid genes, and ∼450 identified metabolites relative to M0 MΦs. Signal protein and expressed gene profiles grouped the MΦs into inflammatory (M1 and M2b) and wound resolution (M2a, M2c, and M2d) phenotypes; however, each had a unique metabolic profile. While both M1 and M2b MΦs shared metabotype profiles consistent with an inflammatory signature; key differences were observed in the TCA cycle, FAO and OXPHOS. Additionally, M2a, M2c and M2d MΦs all profiled as tissue repair MΦs; however, metabotype differences were observed in multiple pathways including hexosamine, polyamine, and fatty acid metabolism. These metabolic and other key functional distinctions suggest phagocytic and proliferative functions for M2a MΦs, and angiogenesis and ECM assembly capabilities for M2b, M2c and M2d MΦs. By integrating metabolomics into a systems analysis of MΦ phenotypes, we provide the most comprehensive map of MΦ diversity to date, along with the global metabolic shifts driving MΦ functional plasticity in these phenotypes.

**Summary Sentence:** Macrophage functional plasticity of six macrophage phenotypes correlates with unique distinctions in cell-surface marker expression, signal protein secretion, transcriptomics profiles, and metabolic processes.

## Introduction

Macrophages (MΦs) display remarkable functional plasticity and the ability to activate diverse responses to a host of intracellular triggers, external stimuli, and nutritional availability. Generally, MΦs are divided into M1(classically activated, pro-inflammatory) MΦs and a broad set of M2 (alternatively activated, anti-inflammatory) MΦs. M2 MΦs have been further subdivided into M2a, M2b, M2c, and M2d subtypes [1]. These M2 subtypes have exhibited numerous, often overlapping, effector functions [2], and have emerged as keystone species in the functional ecology of such diverse environments as wounds [3], tumors [4–6], developing fetuses [7], and systemic autoimmune diseases [8, 9]. Comprehensive, functional characterization of such a diverse array of MΦ populations presents an immense challenge and requires utilization of a systems-based approach that integrates transcriptional, proteomic, and metabolomic profiles to fully characterize these diverse MΦ phenotypes. The emergent field of metabolomics has established a pivotal role for metabolism in immunomodulation of MΦ polarization and functional phenotypes (reviewed in O-Neill, et al. [10]); yet the majority of this research is limited to comparing the more established M1 and M2a phenotypes, leaving much to be learned about the other known M2 MΦ subtypes (i.e., M2b, M2c, and M2d).

Classically activated, pro-inflammatory M1 MΦs are polarized by exposure to IFN-γ, TNF, LPS, and/or other antimicrobial signals that trigger TLR ligation [11]. M1 MΦs exhibit increased and sustained pro-inflammatory responses including production of such anti-microbial molecules RNS and ROS, secretion of inflammatory mediators including IL-6, IL-1β, IL-12, IL-18, and TNFα [12]. The upregulation of MHC-II and T-cell priming surface markers (e.g., CD40, CD80, and CD86), as well as chemokine ligand production (e.g., CXCL9, CXCL10, and CXCL11) in M1 MΦs promote Type I immunity through Th1 and Th17 cytotoxic adaptive immune responses [12]. Although essential for defense against infection, continuous M1 polarization of MΦs can lead to impaired wound healing and significant collateral tissue damage from sustained inflammation. Phagocytosis of apoptotic cells, detection of markers known as DAMPs or alarmins, and production of IL-5, IL-4, and IL-13 by nearby cells dampens M1-driven inflammation and facilitates the onset of an anti-inflammatory, tissue repair response dominated by M2 MΦ functional phenotypes [13].

M2 MΦs (i.e., M2a, M2b, M2c, and M2d) known to represent the opposite end of the polarization spectrum from M1 MΦs, exhibit tissue-reparative, pro-fibrotic, pro-angiogenic, and phagocytic functions [14–16]. Associated with Type 2 immunity and Th cell priming, M2 MΦs are proposed to counteract the inflammatory microenvironment created by the M1 MΦ phenotype [14]. M2a MΦs, polarized by exposure to IL-4 and IL-13, have been described as expressing high levels of MHC-II, MRC1, CD36, and IL-1Ra to promote Th cell activation and suppress IL-1 mediated inflammation [14, 17]. Additionally, M2a MΦs have been shown to secrete chemokines, fibronectins, and growth factors (e.g. FGF and TGF-β) to initiate cellular proliferation, recruit tissue-repair cellular populations, promote angiogenesis, and support development of myofibroblasts [18].

M2b MΦs, polarized by exposure to LPS and immune complexes (ICs), have been shown to crosslink Fcγ receptors via ICs, inducing elevated CD80/CD86 expression for T-cell priming, promotion of Type 2 immunity, and high IL-10 and low IL-12 secretion [14, 19, 20]. Observed to be ineffective at bacterial clearance [21], M2b MΦs display characteristics indicative of a tissue remodeling functional phenotype which include ECM synthesis and angiogenic potential [22].

Resting MΦ exposure to IL-10, glucocorticoids, or TGF-β, drives polarization towards the M2c MΦ, and has been characterized as having increased expression of the *MerTK*, surface display of CD163, secretion of PGF, and activation of MMPs. Based on this profile, M2c MΦs are proposed to promote tissue repair through efferocytosis, ECM remodeling, and angiogenesis [15, 23]. While this phenotype appears to overlap with the M2a MΦ functional phenotype, Gabriel, et al. (2014) demonstrated differences in the timing of gene expression during polarization and proposed that this temporal polarization may distinguish these MΦ phenotypes *in vivo* [24]. Additionally, Jetten, et al. (2014) showed that while PGF signaling was necessary for M2c MΦ-induced angiogenesis, PGF was not required for angiogenesis promoted by M2a MΦs [18].

Tumor associated MΦs (TAMs, referred to herein as M2d MΦs), previously investigated within the tumor microenvironment, can be polarized through a variety of mechanisms including co-culture with cancer cells or ascites fluid, exposure to IL-6, LIF, and/or the purine nucleoside adenosine [16]. Associated with potent immunosuppressive functions and the promotion of angiogenesis [16, 25], M2d MΦs have also been shown to secrete numerous proteolytic enzymes (e.g., MMP-2, MMP-9, and cathepsins), growth factors (e.g. EGF, VEGF. PDGF, and bFGF), and anti-inflammatory mediators (e.g., IL-10, TGF-β, B7-H1, B7-H3/H4) [4, 6, 16, 26]. Although the function of M2d MΦs clearly overlaps with other M2 MΦs, the specific role of M2d MΦs within the healing wound remains unknown.

While MΦ phenotypes have yet to be comprehensively described, the plasticity for M1/M2a MΦ has been demonstrated to depend on metabolic immunomodulation [10]. M1 MΦ polarization favors aerobic glycolysis and glycolytic flux into the oxidative branch of the PPP, generating NADPH to feed the oxidative burst and serve as the building block for rapid biosynthesis [10]. Activation of PPP also supports the production of glutathione and other antioxidants to promote cellular longevity under the harsh conditions created by this pro-inflammatory MΦ phenotype [27]. This anti-microbial function is further enhanced by a decoupled anaplerotic TCA with breaks at citrate and succinate. Accumulated citrate from the broken TCA cycle is diverted for the biosynthesis of fatty acids (FAs), prostaglandins, the anti-microbial metabolite itaconate, and the generation of NO [27–29]. Elevated itaconate inhibits SDH, resulting in succinate accumulation, *HIF* stabilization, and *HIF*- mediated expression of pro-inflammatory cytokines [27, 30, 31].

In contrast, M2a MΦs utilize glycolysis to feed an intact TCA cycle coupled with FA oxidation (FAO), and active OXPHOS to generate ATP [32, 33]. FAO also plays a role in reducing inflammation through reduced lipid accumulation and inhibition of pro-inflammatory mediators [34]. Additionally, M2a MΦs rely heavily on glutamine to promote the hexosamine pathway needed for production of N-glycans and the upregulation of the mannose receptor [28]. Finally, arginine flux through ornithine contributes to collagen production and polyamine synthesis [28, 35].

Much less is known regarding metabolic immunomodulation of the remaining M2 MΦ phenotypes (i.e., M2b, M2c, and M2d). Rodriguez-Prados, et al. (2010) demonstrated that much like M1 and M2a MΦs, M2c MΦs utilized glycolysis; however, only M1 MΦs consumed glucose to produce lactate, indicating M1 MΦ use of aerobic glycolysis is unique and both M2a and M2c MΦs utilize glycolysis to drive an intact TCA [30]. In contrast, M2a and M2c MΦs could be distinguished based on glutamine consumption, suggesting a metabolic contribution to the unique mannose expression on M2a MΦs [28]. Elevated glycolysis has also been demonstrated in M2d MΦs, but as a means to promote angiogenesis and tumor metastasis [4]. Like M1 MΦs, M2b cells have been shown to generate NO combined with decreased production of urea [14]. Alternately activated M2 MΦs have an intact TCA, increased OXPHOS, low cellular energy (e.g. high AMP/ATP ratios), and elevated catabolism of FAs [36]; however the metabolic profiles of murine-derived MΦs compared to human-derived MΦs are inconsistent demonstrating how much is still unknown about how metabolism drives function in these diverse MΦ phenotypes [37].

Multiple MΦ functional phenotypes have been described in the literature (reviewed in Anders, et. al., 2019 [38]), yet the key components of metabolic immunomodulation that drive this functional diversity have not been well characterized. With a systems biology approach, we comprehensively examined an *ex vivo* MΦ model that produced six functional phenotypes. Briefly, CD14+ PBMCs isolated from healthy human donors were differentiated into resting MΦs (referred to as M0 MΦs). MΦ subsets were then polarized into M1 (IFN-γ/LPS), M2a (IL-4/IL-13), M2b (IC/LPS), M2c (IL-10), and M2d (IL-6/LIF) MΦ phenotypes. The MΦs were then profiled using a bioanalyte matrix composed of four cell surface markers, ∼50 secreted cytokines/chemokines/growth-factors, ∼800 expressed myeloid genes and ∼450 identified metabolites. By integrating metabolomics into a systems analysis of MΦ phenotypes, we provide the most comprehensive map of MΦ diversity to date, as well as the global metabolic shifts driving MΦ functional plasticity in these phenotypes.

## Materials and Methods

### Human Subjects, Monocyte Isolation, and MΦ Differentiation and Polarization

Written informed consent was obtained from healthy human subjects (ages 18-60 years old) enrolled in the study approved by the VA Puget Sound institutional review board (IRB). Whole blood (100 mL total) was obtained through venipuncture; 70 mL of blood was drawn into 60-mL syringe tubes pretreated with heparin (Fresenius Kabi USA, LLC; Lake Zurich, Illinois, USA) while 30 mL was collected into BD Vacutainer Serum tubes (Franklin Lakes, NJ, USA). The vacutainer serum tubes were incubated at room temperature for 30-60 minutes before centrifugation at 3500 rpm for 20 minutes and separated serum was collected and stored at -20° C. PBMCs were obtained via Ficoll-paque density centrifugation (Ficoll-Paque Plus, GE Healthcare Bio-Sciences; Uppsala, Sweden) followed by CD14+ monocyte isolation via negative immunomagnetic selection using the Pan Monocyte Isolation Kit (Miltenyi Biotech; Bergisch Gladbach, Germany). Monocyte culture media was made of Roswell Park Memorial Institute (RPMI) 1640 (Corning; Manassa, VA, USA) modified with 1% (v/v) each of minimum essential media (MEM) non-essential amino acids (Gibco; Grand Island, NY, USA), sodium pyruvate (Gibco; Grand Island, NY, USA), L-glutamine (Lonza; Walkersville, MD, USA) and supplemented with 10% (v/v) autologous human serum (AHS; pooled from a minimum of four healthy human donors) and 50 ng/mL of recombinant human MΦ colony-stimulating factor (M-CSF) (Biolegend; San Diego, CA, USA). Monocytes were cultured in monocyte culture media at a cellular density of 1.5 x 10^6^ cells/mL in 6-well cell culture plates. and incubated at 37° C with 5% CO_2_. Monocytes received fresh modified culture media on day 4 and day 7 and were treated with predefined polarization stimuli to achieve the desired MΦ phenotype. Cells exposed to M-CSF alone produce the M0 phenotype. Classically activated M1 phenotype was achieved by exposure to 100 ng/mL LPS from *Escherichia coli* O111:B4 (Sigma Aldrich; St. Louis, MO, USA) and 20 ng/mL recombinant human IFN-γ (PeproTech; Rock Hill, NJ, USA). Alternately M2 MΦs were activated as follows: M2a MΦs from stimulation with 20 ng/mL recombinant human IL -4 (PeproTech; Rock Hill, NJ, USA) and 25 ng/mL recombinant human IL-13 (PeproTech; Rock Hill, NJ, USA); M2b cells with 100 ng/mL LPS and 16.7 µL/mL immune complexes (IC; see details below); M2c cells with 25 ng/mL recombinant human IL-10 (PeproTech; Rock Hill, NJ, USA); and M2d MΦs from stimulation with and 50 ng/mL recombinant human IL-6 (R&D Systems; Minneapolis, MN, USA) and 25 ng/mL recombinant human leukemia inhibitory factor (LIF; R&D Systems; Minneapolis, MN, USA). ICs for M2b polarization were generated by mixing 1 µL of a 1.32 mg/mL solution of bovine serum albumin (BSA; MP Biomedicals; Auckland, New Zealand) with 50 µL anti-albumin (bovine serum), rabbit IgG fraction polyclonal antibodies (Invitrogen; Eugene, OR, USA) and incubating at 37° C for one hour. This IC mixture, stable at 4° C for several days, was stored until needed. Polarized cells were incubated at 37° C with 5% CO_2_ from between 24 and 72 hours in accordance with established MΦ differentiation protocols.

### Characterization of MΦ Phenotypes

#### Flow Cytometry Analysis

Flow cytometry was conducted to access differences in cell surface marker expression between the six different phenotypes. Preliminary experiments were conducted using CD14+ monocytes on 2-, 5- and 7-days post-plating to differentiate between monocytes, immature MΦs, and mature M0 MΦs and to identify gating regions for future MΦ phenotype experiments. For MΦ phenotype experiments, a panel of four fluorescent markers was employed including fluorescein isothiocyanate (FITC) mouse anti-human CD40 (BD Biosciences; San Jose, CA, USA), phycoerythrin (PE) mouse anti-human CD86 (BD Biosciences; San Jose, CA, USA), peridinin chlorophyll protein complex cyanine (PerCP-Cy™5.5) mouse anti-human CD163 (BD Biosciences; San Jose, CA, USA) and allophycocyanin (APC) mouse anti-human human leukocyte antigen – DR isotype (HLA-DR) (BD Biosciences; San Jose, CA, USA). Unstained cells, single stained cells, and fluorescent minus one (FMO) controls were included for each analyzed phenotype to identify unstained cellular populations. All samples were evaluated on an Accuri C6 flow cytometer (BD Biosciences; San Jose, CA, USA) and the data analyzed using Kaluza 2.1 Analysis software (Beckman Coulter; Indianapolis, Indiana, USA).

#### Intracellular Metabolite Isolation and Analysis

With the goal of creating a comprehensive snapshot of cellular function, multiple experimental sample types were collected from each MΦ phenotype during cell harvest to obtain transcriptomic, proteomic, and metabolomic data from each experimental trial. At the desired time point, the cellular supernatant, composed of cell culture media and any non-adherent cells, was removed from the cell culture well and placed into falcon tubes on ice. One mL of ice-cold phosphate buffered saline (PBS) was added to the adherent cell layers and then removed to the tubes containing the cellular supernatant and the tubes centrifuged at 2000 rpm for 8 minutes. The collected supernatant was stored at -80° C until further analysis. Pelleted cells were flash-frozen in liquid nitrogen and stored on ice, while the remaining adherent cells were quenched with 350 µL of ice-cold 100% methanol (Honeywell; Muskegon, MI, USA) and 350 µL of ice-cold ultrapure distilled water (Invitrogen; Grand Island NY, USA) and removed through gentle cell scraping and added to the cell pellet fraction. Following the addition of 700 µL of ice-cold chloroform (Acros Organics; Thermo Fisher Scientific; Waltham, MA, USA), the collected cells were vortexed for 30 seconds and were transferred to FastPrep® lysing matrix D tubes (MP Biomedicals; Auckland, New Zealand). To achieve cell lysis, the tubes were homogenized during two cycles of 40 s each at 4.0 m/s with a 90 s delay between cycles utilizing the FastPrep-24™ 5G Homogenizer (MP Biomedicals; Auckland, New Zealand). The homogenized samples were centrifuged at 16,000 x g for 5 minutes at 4° C, and then placed immediately on ice. The polar (methanol/water) layer and non-polar (chloroform) layers were subsequently transferred to 1.5 mL protein low binding microcentrifuge tubes. These metabolite suspensions were lyophilized overnight without heat on a Thermo Scientific™ Savant™ ISS110 SpeedVac™ (Waltham, MA, USA) and stored at -80°C until the samples were shipped to Metabolon for further analysis. The remaining interphase layer was flash-frozen in liquid nitrogen and stored at -80°C until RNA extraction.

#### Cytokines, Chemokines and Growth Factors

All cytokines, chemokines and growth factors were measured using the supernatant collected during the intracellular metabolite extraction at 24- and 72-hr post polarization. The analytes were evaluated utilizing multiplex magnetic bead-based immunoassays (ProcartaPlex™, Thermo Fisher Scientific; Waltham, MA, USA) per the manufacturer’s instructions, and were run on a MAGPIX® instrument (Luminex Corporation; Austin, TX, USA).

#### RNA Extraction

RNA was purified utilizing the Qiagen RNAeasy® plus mini kit (Germantown, MD, USA). The lysing tubes containing the interphase layer from the metabolite were first thawed, if needed, on ice, resuspended in the RLT lysis buffer supplemented with 1% (v/v) of β-mercaptoethanol and homogenized, as described above. Following lysis, the RNA was extracted and purified according to the manufacturer’s instructions. The integrity of the extracted RNA was evaluated on an Agilent 2200 Tape Station (Santa Clara, CA, USA) and diluted to a final volume of 5 µl, at a concentration of 20 ng/µL.

#### Nanostring analysis

The nCounter Gene Expression – Hs Myeloid v2 Panel CodeSet (NanoString Technologies; Seattle, WA, USA) was used to evaluate the expression of genes associated with myeloid innate immune system response. In brief, and according to NanoString’s procedure, 100ng of each RNA sample was added to the CodeSet in hybridization buffer and incubated at 65°C for 16 hours. Purification and binding of the hybridized complexes were then carried out automatically on the nCounter Prep Station using magnetic beads derivatized with short nucleic acid sequences that are complementary to the capture and reporter probes. The hybridization mixture was first allowed to bind to the magnetic beads via the capture probe then washed to remove excess reporter probes and unhybridized DNA fragments. Probe complexes were then eluted off the beads and hybridized to magnetic beads complementary to the reporter probe followed by wash steps to remove excess capture probes.

In the last step, the purified target-probe complexes were eluted off and immobilized in the cartridge for data collection which was carried out in the MAX System Digital Analyzer (NanoString Technologies; Seattle, WA, USA). Digital images were processed, and barcode counts tabulated in a comma-separated value (CSV) format. The raw count data was then normalized to the mean of positive control probes followed by RNA content normalization to the geometric mean of housekeeping genes in the CodeSet. Data normalization was performed using nCounter Analysis Software version 4.0 (NanoString Technologies; Seattle, WA, USA). Gene set enrichment analysis (GSEA) was utilized to determine which Nanostring-defined functional pathways were regulated within the individually polarized phenotypes. Two representations of these pathways are presented in the data. The first, undirected global significance (UGS) score, was utilized to identify which pathways were differentially regulated in each phenotype regardless of whether the genes were up- or down-regulated when compared to the M0 MΦs. An extension of this score, the directed global significance statistics (DGS), measured the extent to which the genes were up- or down-regulated within the given pathway. In general, a positive pathway score for a specific phenotype indicates a predominance of genes that are up-regulated within the pathway as compared to those that are down-regulated; a negative pathway score indicates the reverse [39]. Additional analyses were conducted using bioDBnet Biological Database [40] and DAVID bioinformatics resources [41]. The top up-regulated genes within each activated phenotype were selected for gene ontology (GO) enrichment analysis. The GO processes significantly enriched (p ≤ 0.05) were then chosen for comparison. With further evaluation, processes common to all phenotypes and related to either specific diseases or ubiquitous to the immune systems were eliminated from the GO process list (e.g. GO:0045824: negative regulation of innate immune response).

#### Metabolite Analysis

All samples were analyzed by Metabolon (Morrisville, NC, USA) using four ultra-high-performance liquid chromatography/tandem accurate mass spectrometry (UHPLC/MS/MS) methods. The following is a summary of Metabolon’s procedure. All methods utilized a Waters ACQUITY ultra-performance liquid chromatography (UPLC) and a Thermo Scientific Q-Exactive high resolution/accurate mass spectrometer interfaced with a heated electrospray ionization (HESI-II) source and Orbitrap mass analyzer operated at 35,000 mass resolution. Lyophilized samples were reconstituted in solvents compatible to each of the four methods and contained a series of standards at fixed concentrations to ensure injection and chromatographic consistency. One aliquot was analyzed using acidic positive ion conditions, chromatographically optimized for more hydrophilic compounds. In this method, the extract was gradient eluted from a C18 column (Waters UPLC BEH C18- 2.1×100 mm, 1.7 µm) using water and methanol, containing 0.05% perfluoropentanoic acid (PFPA) and 0.1% formic acid. A second aliquot was also analyzed using acidic positive ion conditions that was chromatographically optimized for more hydrophobic compounds. In this method, the extract was gradient eluted from the same aforementioned C18 column using methanol, acetonitrile, water, 0.05% PFPA and 0.01% formic acid and was operated at an overall higher organic content. A third aliquot was analyzed using basic negative ion optimized conditions using a separate dedicated C18 column. The basic extracts were gradient eluted from the column using methanol and water, however with 6.5 mM ammonium bicarbonate at pH 8. The fourth and final aliquot was analyzed via negative ionization following elution from a HILIC column (Waters UPLC BEH Amide 2.1×150 mm, 1.7 µm) using a gradient consisting of water and acetonitrile with 10mM ammonium formate, pH 10.8. The MS analysis alternated between MS and data-dependent MS scans using dynamic exclusion. The scan range varied slighted between methods but covered 70-1000 m/z. Several types of controls were analyzed in concert with the experimental samples: a pooled matrix sample generated by taking a small volume of each experimental sample served as a technical replicate throughout the data set; extracted water samples served as process blanks; and a cocktail of quality control (QC) standards that were carefully chosen not to interfere with the measurement of endogenous compounds were spiked into every analyzed sample, allowed instrument performance monitoring and aided chromatographic alignment. Instrument variability was determined by calculating the median relative standard deviation (RSD) for the standards that were added to each sample prior to injection into the mass spectrometers. Overall process variability was determined by calculating the median RSD for all endogenous metabolites (i.e., non-instrument standards) present in 100% of the pooled matrix samples. Experimental samples were randomized across the platform run with QC samples spaced evenly among the injections.

Raw data was extracted, peak-identified and QC processed using Metabolon’s hardware and software. Compounds were identified by comparison to library entries of purified standards or recurrent unknown entities. Biochemical identifications are based on three criteria: retention index within a narrow RI window of the proposed identification, accurate mass match to the library +/- 10 ppm, and the MS/MS forward and reverse scores between the experimental data and authentic standards. The MS/MS scores are based on a comparison of the ions present in the experimental spectrum to the ions present in the library spectrum. Peaks were quantified using area-under-the-curve. The resulting data was then normalized to cell count to account for differences in metabolite levels due to differences in the amount of material present in each sample.

#### Metabolomics Data Analysis

After normalization, peak intensity values were uploaded into MetaboAnalyst 4.0 (Ste. Anne de Bellevue, Quebec) [42], normalized using pareto scaling, and then statistically analyzed. Using MetaboAnalyst 4.0, partial least-squares to latent structures discriminant analysis (PLSDA) and orthogonal PLSDA (OPLSDA) were employed to characterize and visual metabolomic differences between the resting MO MΦs and the individually polarized phenotypes. For detailed descriptions of OPLSDA utilization with metabolomics data, readers are referred to Wiklund, et al., (2008) [43]. PLSDA, a regression method, was employed to identify relationships between the UHPLC/MS/MS data (X) and binary vectors (Y) with a value of 0 for M0 resting MΦs and 1 representing the polarized phenotype as they are compared individually to the M0 cells. Orthogonal PLS (OPLS) was then used to separate the systematic variation in X into two components: the first component, a predictive component (covariance), linearly associated with Y, describes between phenotypic variance; and the second component (correlation), orthogonal to Y describes between sample variation [43].

To adequately demonstrate the OPLSDA results, three data visualization methods were utilized. The first, cross-validated score plots, display the cross-validated score values (Orthogonal T-score) on the y-axis versus the predictive or model T-score (T-score) on the x-axis for each sample point within the analysis. S-Plots were used to visualize the predictive component of the model. The x-axis represents the magnitude or covariance of each metabolite to the model while the reliability or correlation of those metabolites is plotted on the y-axis. Generally, metabolites that have covariance scores greater than 0.2 or less than -0.2. and correlation values greater than 0.6 or less than -0.6 are considered to contribute significantly to the predictive value of the model. Shared and Unique Structure (SUS) plots were employed to compare the results from the M1 OPLS model to each M2 (M2a, M2b, M2c and, M2d) phenotype model. For example, by plotting the correlation from the predictive component from the M0 versus M1 model and the M0 versus M2a model, metabolic biomarkers can be identified that are not only uniquely correlated to the M1 and M2a MΦs, but also have shared differentiation from the parent M0 phenotype. The correlation scores for each M2 subtype were plotted on the y-axis and the M1 scores on the x-axis. In this representation, metabolite features that are shared between phenotypes will cluster close to the diagonals while those features unique to each phenotype will fall outside of the diagonals.

### Online Supplemental Material

Additional data provided in the Online Supplemental Material includes gene and protein expression of immunomodulatory factors (Supplemental Figure 1). Supplemental Figure 2 details global transcriptomics differences between each MΦ functional phenotype and the parent M0 phenotype. Supplemental Figure 3 includes representations of PCA and OSPLDA multivariate analysis of the global transcriptomic data and a heatmap demonstrating metabolic differences between each MΦ functional phenotypes relative to the parent, resting macrophage phenotype. Additional phenotypic data for the for γ-glutamyl amino acids and various lipid subgroups are depicted in Supplemental Figure S4. Supplemental Tables S1 and S2 outline the relative genetic expression of select transcripts and the relative concentrations of select pathway metabolites for each polarized phenotype compared to M0 parent cells, respectively.

## Results

### Classification of MΦ Phenotypes by Cell Surface Markers Provides Poor Resolution of Subtypes

Six MΦ phenotypic subpopulations were produced from CD14+ monocytes derived from human PBMCs by differentiated into resting MΦs (referred to herein as M0 MΦs) prior to polarization into M1 (IFN-γ/LPS), M2a (IL-4/IL-13), M2b (IC/LPS), M2c (IL-10), and M2d (IL-6/LIF) MΦ phenotypes. For simplicity of notation, these phenotypes will be referenced herein as M1, M2a, M2b, M2c, and M2d MΦs. Following polarization, these MΦ phenotypes were comprehensively profiled by cell-surface marker display, immunity-targeted proteomics, myeloid gene transcriptomics, and global metabolomics to obtain a systems-level functional profile for each phenotype.

Four MΦ cell surface markers commonly associated with antigen-presentation or phagocytotic capacity of MΦ phenotypes [44] were used as a first-pass characterization of phenotype function, namely CD40, CD86, CD163, and HLA-DR (Fig. 1). As shown in the histogram overlay of phenotype (Fig. 1 A-D) and associated MFI bar chart (Fig. 1E-H), pro-inflammatory M1 MΦs (red bars) demonstrated elevated cell-surface display for CD40 (Fig 1A, E) and CD86 (Fig. 1B, F) compared to resting MΦs (M0, blue bars). Within the alternatively activated MΦ subgroups, only the M2a MΦs (yellow bars) indicated APC functionality with higher display levels of both CD86 (Fig. 1B, F) and HLA-DR (Fig. 1D, H). Cell-surface display of CD163 (Fig. 1C, G) was elevated in both M2c (gray bars) and M2d MΦs (purple bars), while all other phenotypes had decreased cell-surface display when compared to M0 MΦs. Interestingly, M2b MΦs (green bars) alone had decreased cell-surface display of HLA-DR when compared to the M0 parent phenotype (Fig. 1D, H). Taken together, M1 MΦs profiled as CD40^high^CD86^high^CD163^low^, M2a MΦs as CD86^high^CD163^low^HLA-DR^high^, M2b MΦs as CD163^low^HLA-DR^low^, while both M2c and M2d MΦs appeared as CD86^low^CD163^high^. Based on cell-surface display of these markers, APC functionality is primarily exhibited by M1 and M2a MΦs, T-cell stimulation is dominated by M1 MΦs, and phagocytic capacity was predominant in M2c and M2d MΦs. Though common indicators of the broad functionality of MΦs, these cell-surface display profiles clearly lack the resolution to distinguish between these six phenotypes, nor provide much insight into phenotypic functionality.

**Figure 1:**
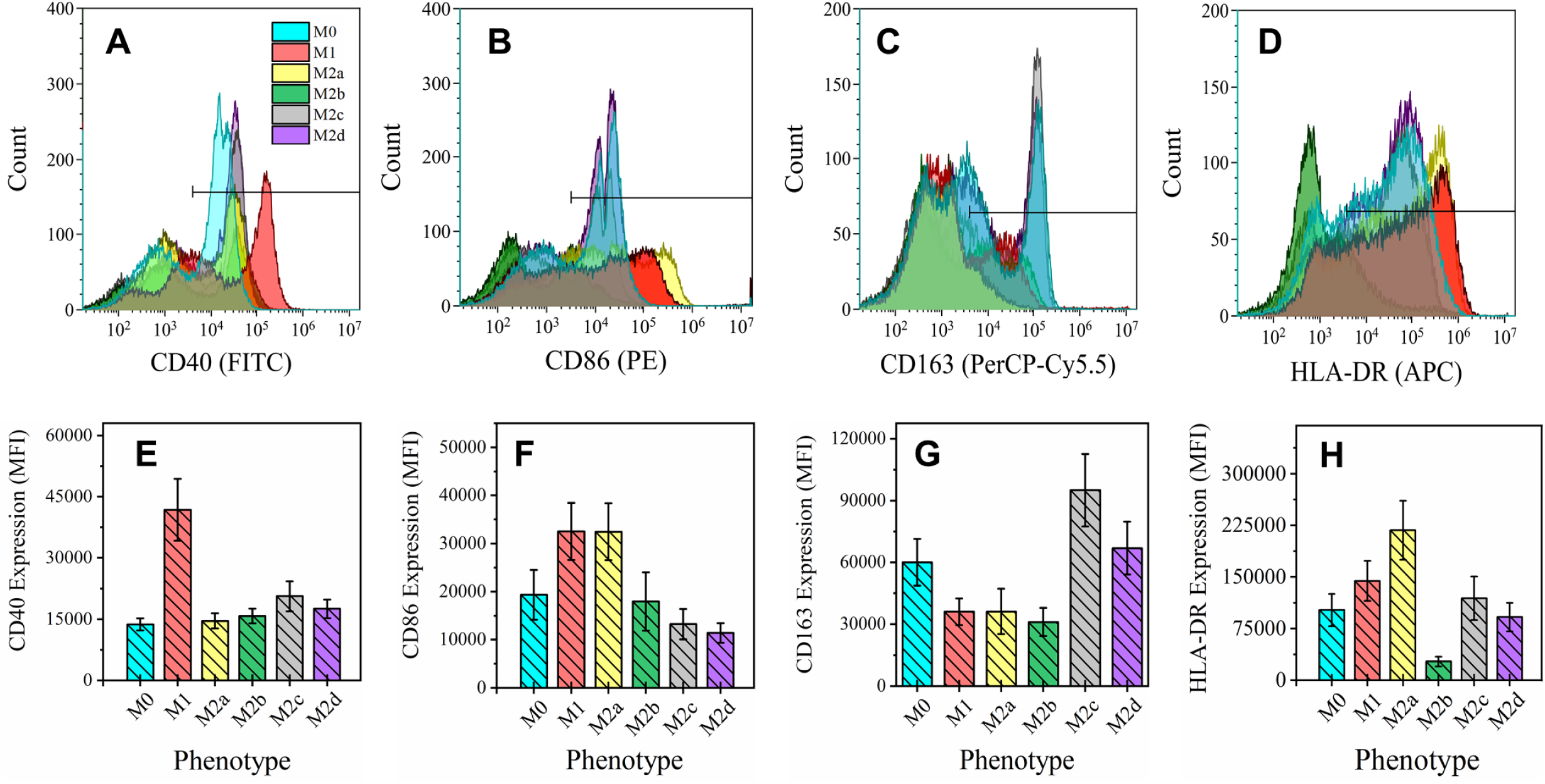
Macrophage Functional Phenotyping by Common Cell-Surface Markers. Cell surface marker profile of six MΦ phenotypes were detected by FACS. CD14+ monocytes were isolated from human blood-derived PBMCs, differentiated into resting MΦ (M0, shown in blue) with M-CSF *ex vivo*, and polarized into five activated phenotypes using IFN-γ/LPS (M1, shown in red), IL-4/IL-13 (M2a, shown in yellow), IC/LPS (M2b, shown in green), IL-10 (M2c, shown in gray), or IL-6/LIF (M2d, shown in purple) for 72 hours. FACS gating on four common MΦ cell surface markers for all six MΦ phenotypes is shown as live cell count for each marker gate (A-D) and mean fluorescence intensity (MFI) profile of gate (E-H). The cell surface markers depicted include CD40 **(A, E)**, CD86 **(B, F)**, CD163 **(C, G)**, and HLA-DR MHC class II receptor **(D, H)**. Histograms are total cell count and representative of three biological replicates. MFI is normalized to total cell count and includes all three biological replicates (N=3, mean ± SEM).

### Targeted Profiling of Immune-Mediator Profile by both Transcript and Protein Expression Demonstrates MΦ Functional Phenotype Along Pro-/Anti-Inflammatory Axis

Cytokines, chemokines, and growth factors induced during MΦ activation are involved in multiple effector functions, including the promotion or inhibition of inflammation. Post-polarization, MΦ culture supernatant was multiplex profiled for 22 immunomodulating proteins as depicted (Fig. 2A-C, bar charts, left y-axis and Supplemental Fig. 1A-G in the Online Supplementary Material). In parallel, total RNA was extracted from each MΦ culture and profiled for mRNA expression (Fig. 2A-C, diamond-whisker plots, right y-axis and Supplemental Fig. 1A-G). As IL-10, IL-4, IL-13, IFNγ, and IL-6 were used as polarizing factors for M2c, M2a, M1, and M2d MΦs, respectively, an N/A was used for these specific phenotypes and respective protein concentration profiles.

**Figure 2:**
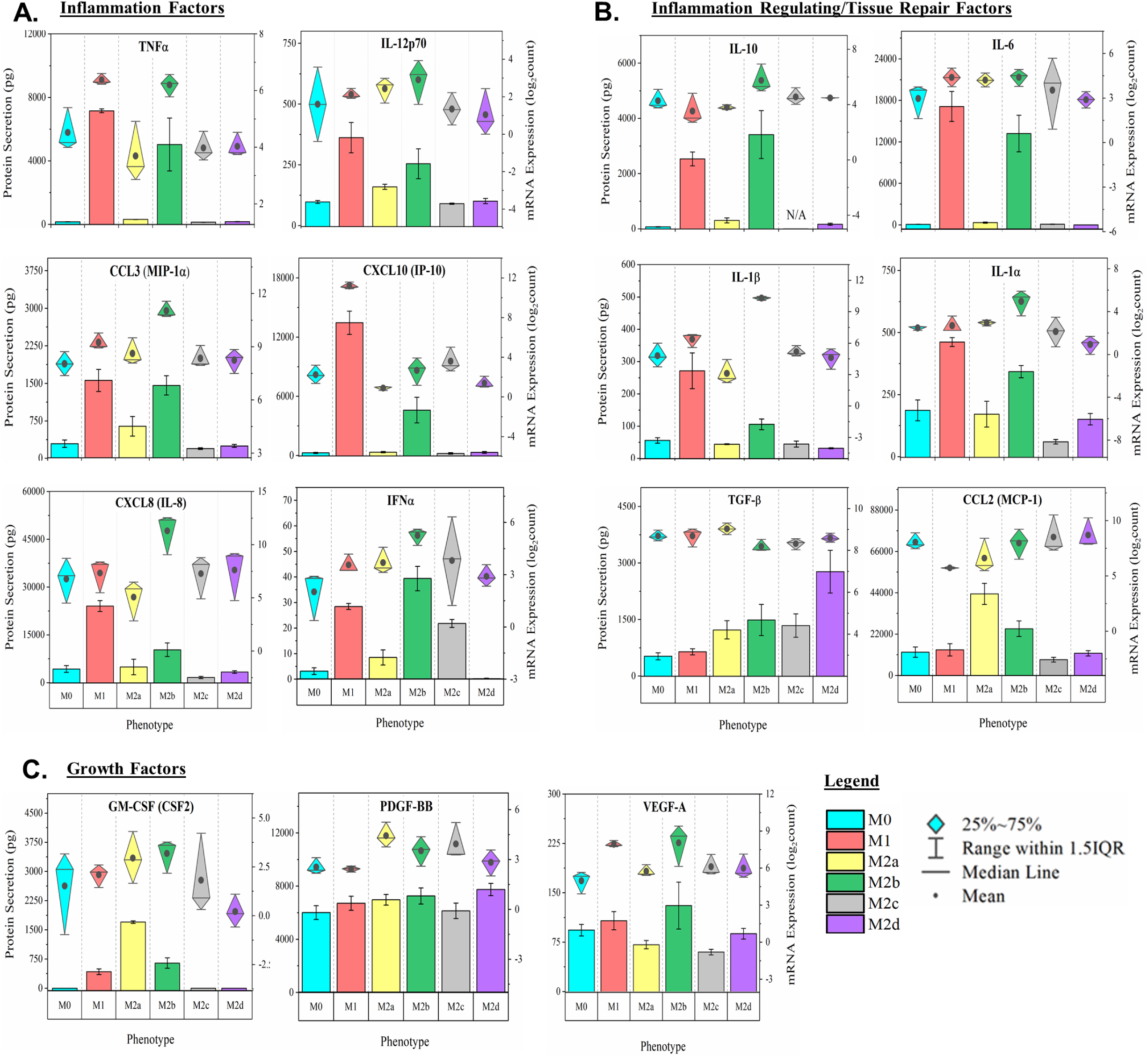
MΦ Functional Phenotyping by Gene and Protein Expression of Immunomodulatory Factors. Multiplex detection of immunomodulatory factors in six MΦ phenotypes harvested at 24 hours were detected using magnetic bead-based quantification of mRNA and secreted protein. CD14+ monocytes were isolated from human blood-derived PBMCs, differentiated into resting MΦ (M0, shown in blue) with M-CSF *ex vivo*, and polarized into five activated phenotypes using IFN-γ/LPS (M1, shown in red), IL-4***/***IL-13 (M2a, shown in yellow), IC/LPS (M2b, shown in green), IL-10 (M2c, shown in gray), or IL-6/LIF (M2d, shown in purple). Gene and protein expression profile (diamond-whisker plots and bar charts, respectively) of key functional molecules were profiled by multiplex assay after 24 hours of *ex vivo* polarization. Immunomodulatory function includes pro-inflammatory (TNF⍺, IL-12p70, CCL3, CXCL10, CXCL8, and IFN⍺) **(A)**, immune-regulatory/tissue-repair (IL-10, IL-6, IL-1b, IL-1⍺, TGF-b, and CCL2) **(B)**, and growth factors (GM-CSF, PDGF-BB, and VEGF-A) (**C)**. Diamond-whisker plots display 25%-75% quartile range, median, and mean. Bar charts indicate mean ± SEM. Expression profiles were normalized to total cell count and include three biological replicates (N=3).

As expected, M1 MΦs secrete high levels of pro-inflammatory cytokines and chemokines including TNFα, IL-12p70, CCL3, CXCL10, CXCL8, and IFNα (Fig. 2A); however, the M2b MΦs, which are often classified under the alternately-activated M2 MΦ phenotypes, also produced these pro-inflammatory factors, though to a lesser extent (Fig. 2A). Additionally, both M1 and M2b MΦs phenotypes also secreted anti-inflammatory cytokines IL-4 (Supplemental Fig. 1D) and IL-13 (Supplemental Fig. 1E), demonstrated to mediate inflammatory responses and drive M2a MΦ activation [45]. Other inflammation-mediating cytokines secreted by both M1 and M2b MΦs include IL-1α and IL-6 (Fig. 2B). While both M1 and M2b MΦ profiles indicate functionality on the pro-inflammatory side of the immune-modulation axis, M2b MΦs are moderately different from M1 MΦs in secretion of IL-10, TGFβ, CCL2, and GM-CSF (Fig. 2B, C).

Of the other alternately activated M2 MΦ phenotypes, M2a MΦs are distinguished by secretion of CCL2 (Fig. 2B) and GM-CSF (Fig. 2C), while both M2a and M2b MΦs produced IFNγ (Supplemental Fig. 1F). While demonstrating similar patterns for most pro-inflammatory mediators, M2c MΦs can be distinguished from M2d MΦs by secreting a surprising level of IFNα while M2d MΦs are exceptional in their high levels of TGFβ secretion (Fig. 2 A, B). Taken together, these protein profiles indicate that M1 and M2b MΦs are significantly more active along the inflammatory axis than any of the other MΦ phenotypes and that phenotypic functionality for M2a, M2c, and M2d is much more restricted with regard to inflammation.

Finally, the mean mRNA expression (represented by the dark circles in the diamond-whisker plots) generally reflect trends observed in secreted protein levels with a few notable exceptions. Most dramatically, IL-1β and CXCL8 transcripts in M2b MΦs are significantly upregulated when compared to the other phenotypes, but secreted protein levels are far less remarkable and significantly lower than the M1 MΦ profile (Fig. 2 A, B). These striking differences indicate an important contribution of post-transcriptional regulation to phenotypic functionality, specifically in the M2b MΦ phenotype.

### Global patterns of myeloid gene expression demonstrate impact of polarization and further define functional phenotype as distinct from the parent, resting MΦ phenotype

Univariant, hierarchical clustering (HC) of gene expression levels revealed the modulatory effect polarization has on these MΦ phenotypes. The most impactful dynamics in gene expression were observed in M1, M2a, and M2b MΦs (Supplemental Fig. 2A). Interestingly, M1 and M2b clustered together, while all other M2 subtypes clustered more closely with their non-polarized parent phenotype M0. Of these M2 subtypes, M2a and M2c MΦs formed unique clusters, while M2d MΦs clustered together with M0 MΦs. To further visualize polarization impacts on gene expression dynamics, the fold change for each gene was calculated relative to the parent M0 MΦ phenotype and relative expression was displayed as the distribution across a logarithmic scale (- 1≤ Log_2_FC ≥1; Supplemental Fig. 2B). In general, the relative number of differentially expressed genes observed for each phenotype mirrors the patterns identified through HC, with M1 and M2b MΦs most transcriptionally active and M2a, M2c, and M2d decreasingly active in that order. Interestingly, the relatively increased transcriptional activity observed in both M1 and M2b MΦs is primarily accounted for in upregulated gene expression when compared to the other M2 MΦ subtypes, for which increased transcriptional activity is fairly evenly distributed between up and down regulated genes, further emphasizing the correlation between polarization and activation specific to M1 and M2b MΦ phenotypes.

ANOVA analysis using an adjusted p-value (false discovery rate [FDR]) of 0.05 revealed 189 SDEGs across all phenotypes and reanalysis by HC demonstrated that, when constrained to SDEG, the M2d phenotype more closely clusters with M2c MΦs than their parent M0 MΦs; however, M2c, M2d, and M0 MΦs cluster separately from M2a MΦs (Supplemental Fig. 2C). Supporting the HC pattern with SDEG constraints, multivariate clustering by PCA project M0, M2a, M2c, and M2d MΦ phenotypes occupying overlapping 3D space, while M1 and M2b MΦ phenotypes share a single dimension and are distinctly different from the other MΦ phenotypes (Supplemental Fig. 3A). Accounting for >90% of the variance in the data within the three components, only 7.4 % of the variance is accounted for in the dimension separating the M1 and M2b MΦ phenotypes, further emphasizing the similar impact of polarization on gene expression patterns in these two phenotypes.

Finally, while PCA provided cluster separation and variance across all six MΦ phenotypes, use of OPLSDA provided clearer class separation of the polarized MΦ phenotypes from their non-polarized, parent phenotype M0 in the cross-validated score plots (Supplemental Fig. 3B-F). Contribution of variance to phenotype cluster separation is plotted along the x-axis and variance within phenotype is plotted along the y-axis. Although variance within phenotype is notable for all phenotypes except M1 MΦs, variance between the parent, non-polarized M0 MΦs and the polarized phenotypes is even more evident. In agreement with the SDEG constrained HC and PCA analyses, between phenotype variance is greatest for M1 MΦs and M2b MΦs, relative to M0 MΦs, accounting for ∼40-45% of the total variance (Supplemental Fig. 3B, D), and nearly twice the percent variance for M2a, M2c, and M2d MΦs at ∼20-30% variance relative to M0 MΦs. As with HC analysis, M2d MΦ profiling is most akin to its parent M0 MΦ phenotype, accounting for only 18.8% of the total variance (Supplemental Fig. 3F). Taken together, these data further demonstrate that when looking beyond the classic pro-/anti-inflammatory axis for defining MΦ phenotype, clear distinctions can be made that suggest phenotype plasticity includes a wide range of functionality in these cell subtypes.

### Pathway Analysis and GO Process Characterization Indicates Biological Functionality Within Each Distinct MΦ Phenotype

Having demonstrated that polarization drove each MΦ phenotype towards distinct programs of myeloid gene expression, gene set enrichment analysis (GSEA) was utilized to determine how transcriptional activation translated into putative myeloid functionality as each phenotype progressed away from the resting, parent M0 MΦ phenotype. Analysis by undirected global significance scores (UGS) displayed all detected pathways differentially impacted relative to the resting M0 parent MΦs (displayed as solid colored plots in the radar graphs in Fig. 3). Consistent with the HC results, M1 and M2b MΦ polarization most dramatically impacted pathways associated with myeloid cell functionality (Fig. 3A, C), while a less significant impact was observed for the M2a, M2c, and M2d MΦs (Fig. 3B, D, E).

**Figure 3:**
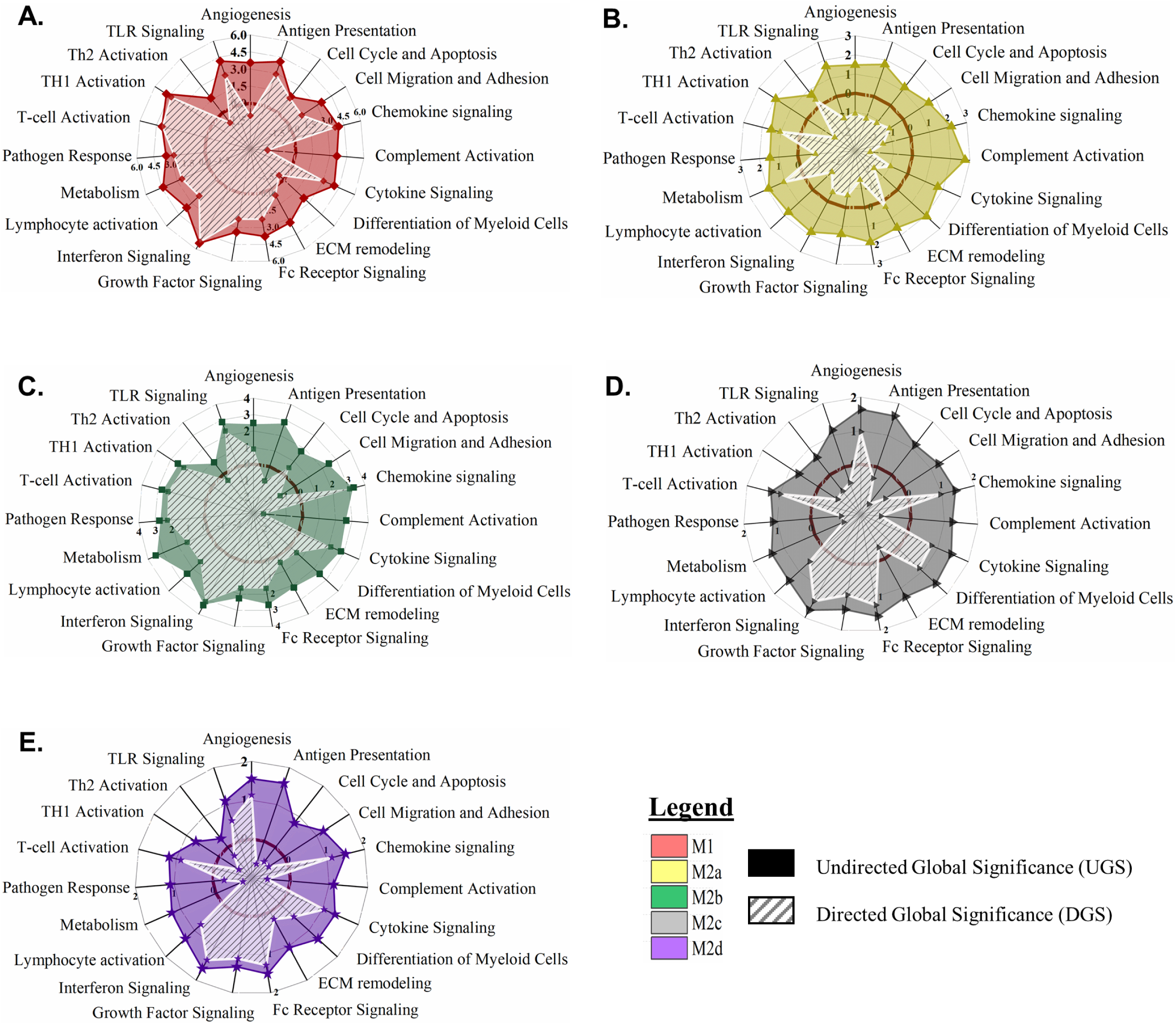
MΦ Polarization into Functional Phenotypes Activates Distinct Gene Expression Profiles Relative to the Parent, Resting MΦ Phenotype (M0). Gene set enrichment analysis (GSEA) and pathway impact scoring of global myeloid gene expression demonstrates distinct patterns in five polarized MΦ phenotypes. Briefly, multiplexed myeloid gene expression was directly detected through molecular barcode probes and normalized to the geometric mean of the housekeeping gene set. GSEA for each phenotype was calculated for pathway impact relative to the non-polarized, resting MΦ (M0). Radar plots for each polarized MΦ phenotype display pathway score based on impact regardless of whether genes were up- or down-regulated (Undirected Global Significance [UGS], solid color) and pathway score based on impact incorporating t-statistic comparison of gene regulation as relatively increased or decreased (Directed Global Significance [DGS], hatch marks). Radar plots include UGS and DGS scores for all five polarized MΦ phenotypes: M1 MΦs (IFN-γ/LPS treated, shown in red) **(A)**, M2a MΦs (IL-4*/*IL-13 treated, shown in yellow) **(B)**, M2b MΦs (IC/LPS treated, shown in green) **(C)**, M2c MΦs (IL-10 treated, shown in gray) **(D)**, and M2d MΦs (IL-6/LIF treated, shown in purple) **(E)**. GSEA based on three experimental replicates for each MΦ s phenotype (N=3).

To better translate the impact of polarization into proposed biological functionality, the directed global significance statistic (DGS) measures the extent to which gene expression contributes to up- or down-regulation of the relevant biological pathway (displayed as white hashed plots in the radar graphs in Fig. 3). Positive DGS values that extend beyond the boundary of impact (thick maroon circle) denote biological pathways up-regulated for that MΦ phenotype, whereas negative DGS values retracted within this boundary denote biological pathways down-regulated for that MΦ phenotype. Highlighted SDEGs of relevance to these biological pathways are depicted in Supplemental Table 1. Of the 19 functionally defined, biological pathways, overall polarization is an “activation” process for M1, M2b, and M2d MΦs with gene expression driving fifteen biological pathways into activation for M1 MΦs and M2b MΦs (Fig. 3 A, C) and eleven biological pathways for M2d MΦs (Fig. 3E). In contrast, M2a MΦ and M2c MΦ gene expression pushed fifteen and ten biological pathways into “inhibition”, respectively (Fig. B, D).

M1 MΦ polarization resulted in gene expression patterns that drove functionality in anticipated directions with up-regulation of antigen presentation, cell migration and adhesion, chemokine signaling, cytokine signaling, interferon signaling, pathogen response, and Th1-specific T-cell activation pathways, while angiogenesis, complement activation, and Th2-specific T-cell activation pathways are downregulated (Fig. 3A). GSEA identified 97 SDEGs ((-1≤ Log_2_FC ≥1 and p ≤ 0.05) contributing to 41% of IFN-γ responsive and 12% LPS-responsive gene expression. Within the IFN-γ responsive genes, interferon pathway genes, *IRF1*, *IRF7*, *IRF8* and *ISG15*, are known transcriptional enhancers of genes associated with antigen presentation and Th1 responses (e.g. *CD4*0 and *CD86*), cytokine and chemokine signaling (e.g. *TNF*, *CXCL9*, *CXCL11*, *CCL4*, *CCL8*, and *STAT1*), and pathogen response (e.g. *IL18* and *IRG1*) [46]. These SDEGs and related upregulated pathways are consistent with defined pro-inflammatory and antimicrobial effector functions known to be associated with M1 MΦ polarization.

While IFN-γ and LPS are considered potent MΦ activators, the IL-4 and IL-13 stimuli utilized for M2a polarization are considered mild polarization activators [47]. Consequently, when compared to the parent M0 MΦ phenotype, M2a MΦs exhibited a remarkably different pathway profile than the M1 MΦs. While all 19 pathways were differentially regulated, only the T-cell activation, T_h_2-specific T-cell activation, metabolism, and ECM remodeling pathways were upregulated (Fig. 3B). Specifically, GSEA results showed that of the 83 identified SDEGs only 36% of these transcripts were upregulated with 84% being downregulated. Despite being upregulated when compared to M0 MΦs, the T-cell and T_h_2 activation pathways did not exhibit any SDEGs using our inclusion criteria. For the ECM remodeling pathway, known metalloproteinase genes, *ADAM19* and *MMP12*, were upregulated and the common M2a MΦ-associated transcripts of *TGM2*, *FABP4*, and *PTGS1* were also upregulated for the metabolism pathway, indicating changes in metabolic processes associated with M2a MΦ polarization. The most upregulated genes, *CCL17*, *CCl22* and *CCL18*, support the generally accepted role that M2a MΦs play in cellular proliferation, ECM remodeling, and tissue repair.

As with our cluster analysis, M2b MΦ proposed biological functionality is unique from all other M2 MΦ subtypes and most like the M1 MΦ phenotype when compared to the resting M0 MΦs. M2b MΦ gene expression during polarization drives downregulation of antigen presentation, cell migration and adhesion, complement activation, and T_h_2-specific T-cell activation response pathways (Fig. 3C). Chemokine signaling, the most upregulated pathway, is associated with prominent SDEGs and is intertwined with many other functional pathways observed for this phenotype. For example, chemokines *CXCL2*, *CXCL3*, and *CXCL5* promote angiogenesis through the recruitment of angiogenic neutrophils and *CCL5, CCL8*, and *CXCL9* support the recruitment and differentiation of myeloid cells necessary for tissue remodeling functions. While M2b MΦ polarization indicates biological functionality comparable to M1 MΦs, distinct differences in such pathways as angiogenesis and cell migration and adhesion indicate that polarization of this phenotype may have a more localized biological relevance.

In a similar trend to M2a MΦs, the pathway scores observed for M2c MΦs imply that IL-10 is also a mild polarizing stimulus; however, patterns of pathway up- and down-regulation are distinctly different. Of the 19 biological pathways detected, nine had DGS scores indicating gene expression driving pathway upregulation compared to the M0 parent MΦs (Fig. 3D). Key pathways included upregulation of angiogenesis, chemokine signaling, cytokine signaling, differentiation of myeloid cells, Fc receptor signaling, growth factor signaling, interferon signaling, lymphocyte activation, and T-cell activation (Fig. 3D). Interestingly, many of the identified SDEGs have broad effects that regulate inflammation (*DUSP1*, *SOCS3*, *S100A8*, and *S100A9*), vascular insult and neutrophil chemotaxis (*S100A8* and *S100A9*), as well as protection against apoptosis and oxidative stress (*IER3*) [15]. Based on these pathway impact profiles, the biological functionality of polarization into this MΦ phenotype may serve to modulate immune response in the transition to resolution.

At first glance, M2c and M2d MΦs appeared to polarize into phenotypes of similar biological functionality; however, polarization of MΦs to the M2c phenotype resulted in a majority of down-regulated pathways and polarization of MΦs to the M2d phenotype resulted in a majority of up-regulated pathways (Fig. 3D, E). Their transcriptional profile suggested overlapping effector functions such as resolution of inflammation (*DUSP1*, *SOCS3*, *THBD*, and *CD163*) and immune suppression (*CCL8*); however, upregulation of TLR signaling in M2d MΦs during polarization is notably distinct from polarization in M2c MΦs and M2d MΦs differentially regulated many known pro-angiogenic markers such as *ENPP2*, *SPHK1*, *CXCL2*, *CXCL3*, and *ID1*, findings in line with a proposed biological role beneficial to survival of associated tumors.

While UGS and DGS scores provided both magnitude and directionality of polarization of each MΦ phenotype overlaid on proposed biological functionality, GO annotation of global myeloid gene expression was utilized to provide a direct comparison between each polarized phenotype (Fig. 4). For each GO annotation, all five MΦ phenotypes are displayed; however, the immediate similarity between M1 and M2b MΦs can be observed when the GO annotation is selected for the top 15 GO processes for M1 MΦs and M2b MΦs. GO annotation generally supported DGS analysis of polarization; however, interesting differences were noted. Specifically, secretion of IL-12, regulation of PDGF production, and response to NO were pathways identified as unique to M1 MΦs (Fig. 4A), while regulation of leukocyte migration, neutrophil extravasation, and connective tissue replacement in inflammatory wound healing were pathways identified as unique to M2b MΦ (Fig. 4C).

**Figure 4:**
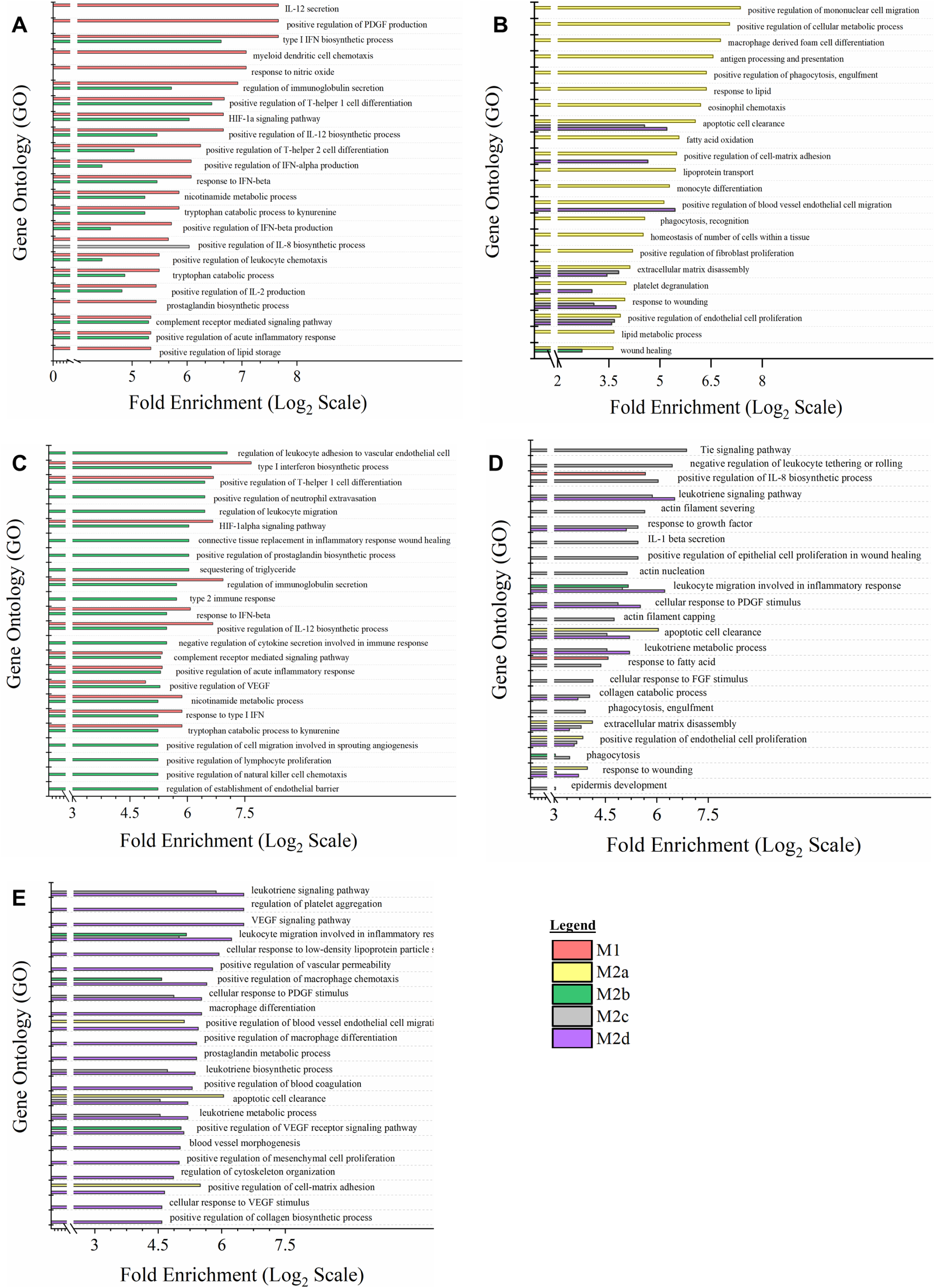
Gene Expression Profile Comparison of Polarized MΦ Phenotypes Identifies Unique and Overlapping Immunomodulatory Functions. Gene Ontology (GO) annotation of global myeloid gene expression in five MΦ phenotypes identifies relative biological functionality across polarized states. Multiplexed myeloid gene expression profile of all five polarized MΦ phenotypes was used to annotate impacted biological functionality based on GO Processes. For each MΦ phenotype, the top 15 impacted GO processes significantly enriched (p ≤ 0.05) were plotted relative to the other four polarized MΦ phenotypes. Enrichment plots for prioritized GO process profile include all five polarized MΦ phenotypes: M1 MΦs (IFN-γ/LPS treated, shown in red) **(A)**, M2a MΦs (IL-4***/***IL-13 treated, shown in yellow) **(B)**, M2b MΦs (IC/LPS treated, shown in green) **(C)**, M2c MΦs (IL-10 treated, shown in gray) **(D)**, and M2d MΦs (IL-6/LIF treated, shown in purple) **(E)**. GO process annotation was based on three experimental replicates for each MΦ phenotype (N=3).

Finally, of the other three M2 MΦ subtypes, GO processes associated with M2a MΦs (Fig. 4B) heavily favored tissue-reparative, pro-fibrotic, pro-angiogenic, and phagocytic functions, while the upregulation of scavengers *CD163* and *MerTk* (Supplemental Table 1) for M2c MΦs are consistent with phagocytosis GO processes detected. Other M2c MΦ identified GO processes included the Tie signaling pathway, known to play a key role in angiogenesis (Fig. 4D). Similarly, for M2d MΦs nine of the top 17 GO processes were causally related to angiogenesis. Of the remaining eight listed processes, six contributed to ECM remodeling (Fig. 4E). In summary, both patterns of polarization detected through GSEA and DGS (Fig. 3) and direct comparison between phenotypes by GO annotation (Fig. 4) demonstrate unique and overlapping proposed biological functionality and further support the reorganization of M2b MΦs from being a M2 subtype to being more closely related to M1 MΦs in functionality.

### Global Metabolomics Demonstrates Functional MΦ Phenotypes are also Distinct Metabotypes

Global metabolomics profile of the parent, resting M0 MΦs and five polarized MΦ phenotypes identified 498 compounds of known identity. The top 50 metabolites, as determined by ANOVA analysis (p < 0.05) and depicted in the heatmap (Supplemental Fig. 3G) demonstrated the immunomodulation of the individual phenotypes. The most striking distinction in the HC analysis is the metabolic differences, especially within lipid metabolism, observed between the M1 MΦs, the M0 MΦs and the M2 subtypes. Of specific interest is the clustering of the M2b MΦs with the other M2 subtypes considering the M2b MΦ propensity towards the M1 pro-inflammatory signature. The notable exception to this trend is the apparent similarities in M1 and M2b MΦ nicotinamide and tryptophan metabolism which is common to pro-inflammatory responses. Additionally, MΦ metabotype clustering was demonstrated through OPLSDA and represented in the cross-validated score plots (inset plots in Fig. 5A.1 - E.1). Separation along the x-axis (between phenotype variation) in the cross-validated score plots quantifies metabotype cluster separation between each polarized MΦ phenotype and the parent M0 MΦs, with cluster separation greatest for M2b MΦs (14.2%) and M1 (13.0%) and the least for M2d (10.3%) and M2c (9.4%) and 8.1% MΦs. The y-axis scores (orthogonal T score or within phenotype variation) reflected the notable variation observed within each MΦ phenotype; however, the variation was similar for each phenotype (∼60%). Furthermore, the larger within phenotype variance observed in this data warrants additional analysis and may indicate uncertainty in the metabotype prediction on the basis of cross-validation scores alone [43].

**Figure 5:**
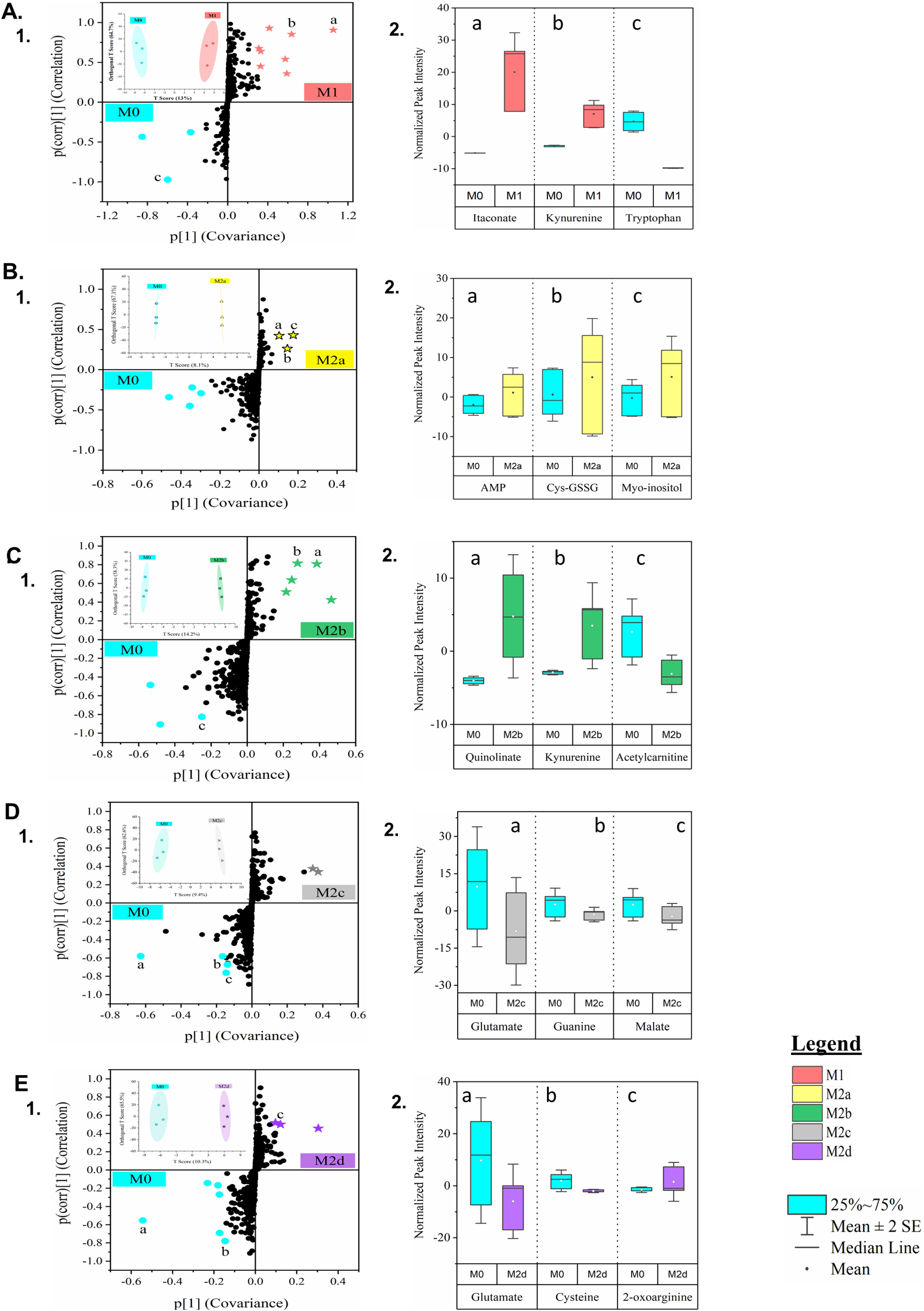
Untargeted Metabolomics of Five Polarized MΦ Phenotypes Identifies Biomarkers of Shared and Unique Profile Relative to the Parent, Resting MΦ Phenotype. Global metabolomics profile of the parent, resting MΦ phenotype (M0) and five polarized MΦ phenotypes identified 498 compounds of known identity, normalized to cell count. MΦ metabotype clustering was determined through Orthogonal Projections to Latent Structures Discriminant Analysis (OPLSDA), inset clustering scores plots with 2D T-scores **(A.1, B.1, C.1, D.1, & E.1)**. S-plot comparison of each polarized MΦ phenotype was plotted relative to the resting, parent MΦ phenotype (denoted in blue) **(A.1, B.1, C.1, D.1, & E.1)** and normalized peak intensity for biomarkers of particular note (a, b, c) are displayed as box-whisker plots **(A.2, B.2, C.2, D.2, & E.2)**. Multivariate, cluster analysis is displayed for all five polarized MΦ phenotypes: M1 MΦs (IFN-γ/LPS treated, shown in red) **(A)**, M2a MΦs (IL-4*/*IL-13 treated, shown in yellow) **(B)**, M2b MΦs (IC/LPS treated, shown in green) **(C)**, M2c MΦs (IL-10 treated, shown in gray) **(D)**, and M2d MΦs (IL-6/LIF treated, shown in purple) **(E)**. Untargeted metabolomics profiles are based on three experimental replicates for each MΦ phenotype (N=3).

To identify putative metabolite biomarkers of high covariance and high correlation separating each polarized phenotype from the parent M0 MΦ phenotype, OPLSDA model contribution from metabolite measurements (covariance; x-axis) and the reliability of the measurements (correlation; y-axis) was projected in S-plots (Fig. 5A.1-E.1). Biomarkers with larger covariance (-0.2 ≤ covariance ≥ 0.2) and correlation (-0.6 ≤ correlation ≥ 0.6) scores can identify biochemically significant metabolites with predictive value for metabolic modeling [43]. Normalized peak intensity for metabolites of interest as putative biomarkers that fall within these parameters are shown for each MΦ phenotype (Fig. 5A.2-E.2), demonstrating the importance of metabolic fluidity to MΦ polarization.

As the generally accepted paradigm is that MΦ phenotypes fall into the classically activated M1 and alternatively activated M2, direct comparison of M1 MΦ metabotype to each M2 MΦ subtype was performed. To achieve this comparison, SUS graphs were employed to visualize OPLSDA characterization of metabolites impacted by polarization (Fig. 6A.1-D.1) and outcomes were projected as both shared and unique metabolite biomarkers between the two metabotypes, as well as associated correlation scores (Fig. 6A.2-D.2). Statistically significant metabolites (FC > 2.0 and p < 0.05) are depicted in the associated volcano plots (Fig. 6A.3-D.3). Contrary to phenotyping by gene expression and correlated functionality, metabotyping each MΦ group did not result in clustering patterns with M2b MΦs more closely related to M1 MΦs and the rest of the M2 subtypes clustered together; however, the unique and shared metabolites within each comparison differed considerably, further indicating the unique metabolism of each phenotype.

**Figure 6:**
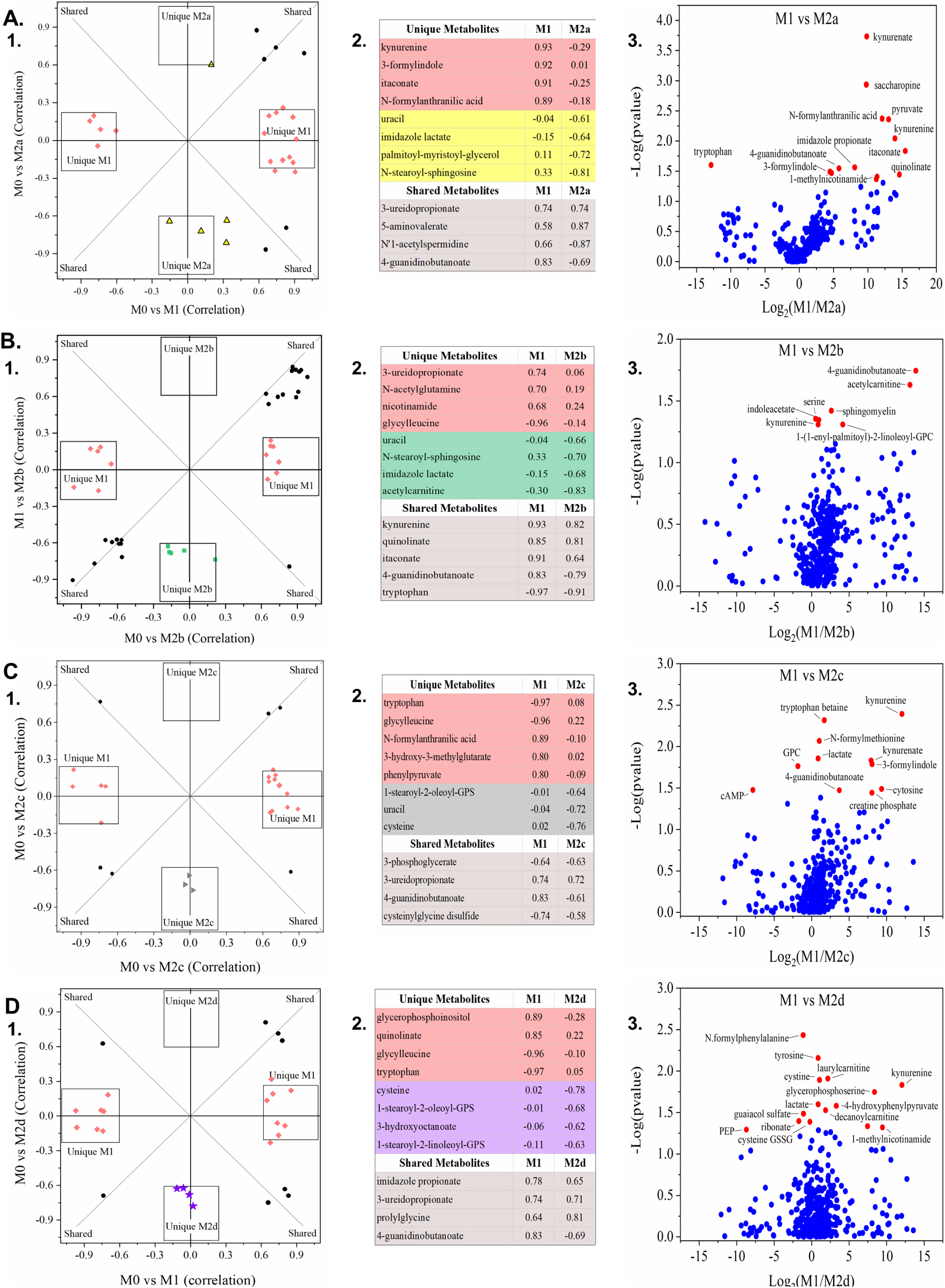
Global Metabolomics Comparison of the Classically Activated MΦ Phenotype (M1 MΦ) to Four Subtypes of Alternately Activated MΦs (M2a, M2b, M2c, & M2d MΦ). OPLSDA clustering relative to the parent, resting MΦ phenotype (M0 MΦ) identified metabolite biomarkers correlated with all five polarized MΦ phenotypes based on OPLSDA variable influence on projection (VIP) values. VIP values were used to identified both shared and unique metabolite biomarkers of the classically-activated M1 MΦ phenotype and four subtypes of the alternately-activated M2 MΦ phenotypes, displayed as OPLSDA Shared and Unique Structures (SUS) plots **(A.1, B.1, C.1, and D.1)**. For each SUS plot, metabolite biomarkers uniquely correlated to the M1 MΦ phenotype (red diamonds) are plotted along the x-axis and those metabolite biomarkers uniquely correlated to the M2 MΦ phenotype are plotted along the y-axis. Shared metabolite biomarkers (black circles) are plotted along the diagonals, reflecting biomarker positive/negative correlation to each polarized MΦ phenotype relative to the parent, resting MΦ phenotype (M0 MΦ). Associated biomarkers and correlation scores are shown in matching tables **(A.2, B.2, C.2, and D.2)**. OPLSDA SUS plots are displayed for four polarized M2 MΦ phenotypes, relative to the M1 MΦs (IFN-γ/LPS treated, shown in red): M2a MΦs (IL-4*/*IL- 13 treated, shown in yellow) **(A)**, M2b MΦs (IC/LPS treated, shown in green) **(B)**, M2c MΦs (IL-10 treated, shown in gray) **(C)**, and M2d MΦs (IL-6/LIF treated, shown in purple) **(D)**. Untargeted metabolomics profiles are based on three experimental replicates for each MΦ phenotype (N=3).

Metabolites impacted by polarization and identified through metabotype comparison between M1 and M2b MΦs (Fig. 5A,C and Fig. 6B) indicated metabolic investment in anti-microbial function with itaconate accumulation (correlation score = 0.91, 0.64, respectively) and antioxidant capacity evidenced by tryptophan consumption (correlation score = -0.97, -0.91, respectively), kynurenine accumulation (correlation score = 0.93), and quinolinate accumulation (correlation score = 0.85, 0.81, respectively), as previously described [48]. Metabolites involved in glutamate metabolism such as N-acetylglutamine (correlation score = 0.70, 0.19, respectively) correlated highly with M1 MΦs only and branched-chain amino acid metabolites such as acetylcarnitine (correlation score = -0.30, -0.83, respectively) were uniquely correlated to M2b MΦs.

As mentioned above, the remaining M2 MΦ subtypes (M2a, M2c, and M2d) cluster separation from the parent M0 MΦ indicated less metabolic divergence upon polarization (Fig. 5B, D, E); however, unique metabolic features could be distinguished between the subtypes. M2a MΦs accumulated cysteine-GSSG and myo-inositol, metabolites involved with glutathione recycling and had similar correlation with uracil as M2b and M2c MΦs, indicating a role for pyrimidine metabolism in these phenotypes (correlation score = -0.61, - 0.66, -0.72, respectively) (Fig. 6A-C). Glutamate was consumed by both M2c and M2d MΦs during polarization and correlation values were also prominent relative to M1 MΦs (correlation score = -0.58, -0.55, respectively) (Fig. 5D-E and data not shown). Arginine catabolism was indicated by M2d MΦ accumulation of 2-oxoarginine (Fig. 5E.2) and for all M2 MΦ subtypes as demonstrated by negative correlation scores for 4- guanidinobutanoate in M2a, M2b, M2c, and M2d MΦ (correlation score = -0.87, -0.79, -0.61, and -0.69, respectively) (Fig. 6A.2-D.2), while histidine metabolism was indicated for M2a and M2b MΦs through negative correlation to imidazole lactate (correlation score = -0.64, -0.68, respectively) (Fig. 6A.2-B.2). Finally, notable differences in lipid metabolism between the M1 and M2 MΦ phenotypes were exhibited by the accumulation of fatty acids and their derivatives in M1 MΦs and corresponding negative correlations within the M2 MΦ subtypes. For example, M2a MΦ negative correlation with palmitoyl-myristoyl-glycerol (correlation score = -0.72), M2a and M2b MΦ negative correlation with N-stearoyl-sphingosine (correlation score = -0.81, - 0.70, respectively), M2c and M2d negative correlation with 1-stearoyl-2-oleoyl-GPS (correlation score = -0.64, -0.68, respectively), and M2d MΦ negative correlation to 1-stearoyl-2-linoleoyl-GPS (correlation score = -0.63) (Fig. 6B.2-D.2; Supplemental Fig. 4B-F).

### Metabolic Pathway Analysis to Defined Unique Metabotypes for Each MΦ Polarized Phenotype

Using topological mapping of global metabolomics organized by p-value from pathway enrichment analysis (y-axis) relative to impact score from pathway topology analysis (x-axis), metabotype was further characterized for each polarized MΦ phenotype and provided a global view of metabolic impact from polarization. For each node, color intensity and size illustrate p-value significance and pathway impact, respectively (Fig. 7). To further support metabotyping by global metabolomics pathway analysis, metabolism-related SDEGs up- or down-regulated relative to the resting parent M0 MΦs are displayed (green or blue, respectively) (Fig. 8). Finally, a comprehensive summary of this metabolic analysis is illustrated in Figure 9. Each colored box above the listed metabolite represents significant fold change relative to the parent M0 MΦs (1 < Log_2_ < -1). Numerical values corresponding to the color scale are included in Supplemental Table 2.

**Figure 7:**
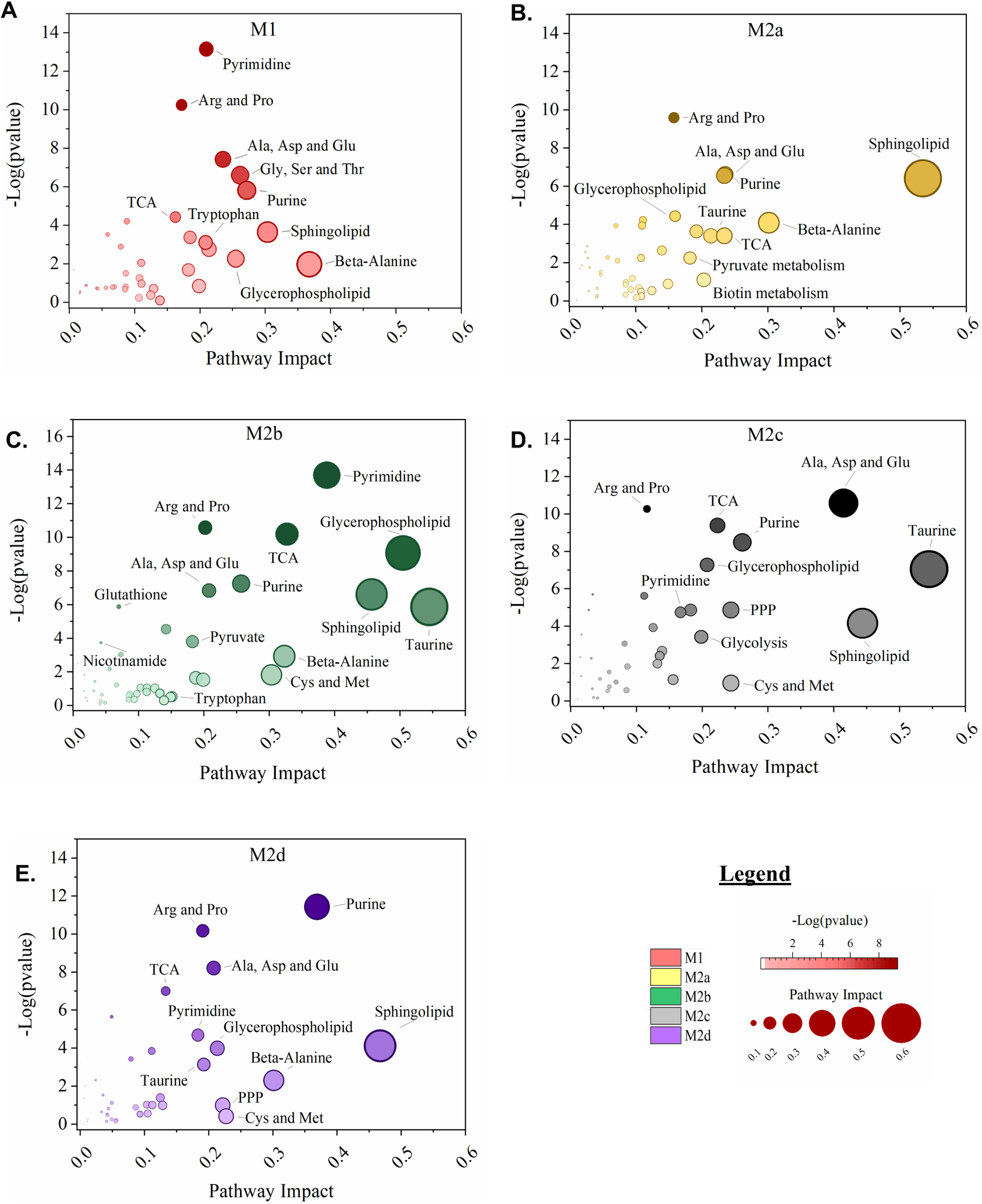
MΦ Polarization into Functional Phenotypes Activates Distinct Metabolic Pathways. Untargeted, global metabolomics profile of polarized MΦ phenotypes identified 498 known compounds, normalized to total cell count, and profiled relative to the resting M0 MΦ phenotype. For normalized, M0-relative metabolite profiles, over-representation analysis by hypergeometric testing and pathway topology analysis by two node centrality measures (degree centrality and betweenness centrality) was performed for each polarized MΦ phenotype including M1 MΦs (IFN-γ/LPS treated, shown in red) **(A)**, M2a MΦs (IL-4***/***IL-13 treated, shown in yellow) **(B)**, M2b MΦs (IC/LPS treated, shown in green) **(C)**, M2c MΦs (IL-10 treated, shown in gray) **(D)**, and M2d MΦs (IL-6/LIF treated, shown in purple) **(E)**. Metabolic Pathway Topology is plotted as Pathway Impact (x-axis) and significance of pathway topography (-Log [p value], y-axis) for each polarized MΦ phenotype.

**Figure 8:**
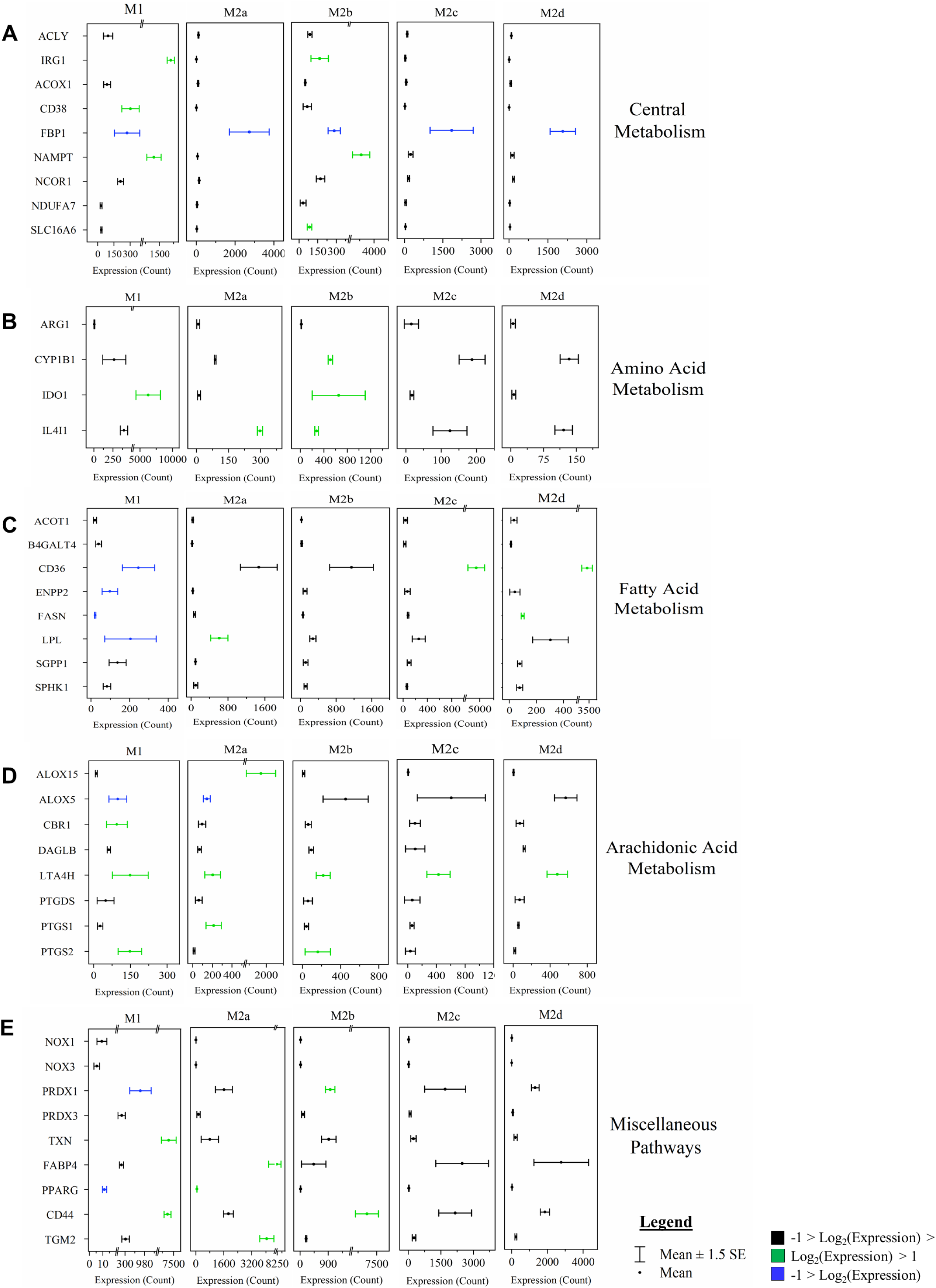
MΦ Polarization into Functional Phenotypes Activates Distinct Metabolic Gene Expression Profiles Relative to the Parent, Resting MΦ Phenotype (M0). Gene set enrichment analysis (GSEA) and pathway impact scoring of global myeloid gene expression demonstrates distinct patterns in metabolism-related transcripts in the five polarized MΦ phenotypes. Associated metabolism gene expression profiles for most impacted pathways are shown including central metabolism **(A)**, amino acid metabolism **(B)**, fatty acid metabolism **(C)**, arachidonic acid metabolism **(D)**, and miscellaneous pathways **(E)**. Red bars represent gene targets that are differentially regulated (p ≤ 0.05) when compared to the M0 phenotype. Myeloid gene expression was directly detected through molecular barcode probes and normalized to the geometric mean of the housekeeping gene set. Gene expression is mean ± SEM. Pathway topology and metabolic gene expression profiles are based on triplicate, experimental replicates (N=3).

**Figure 9:**
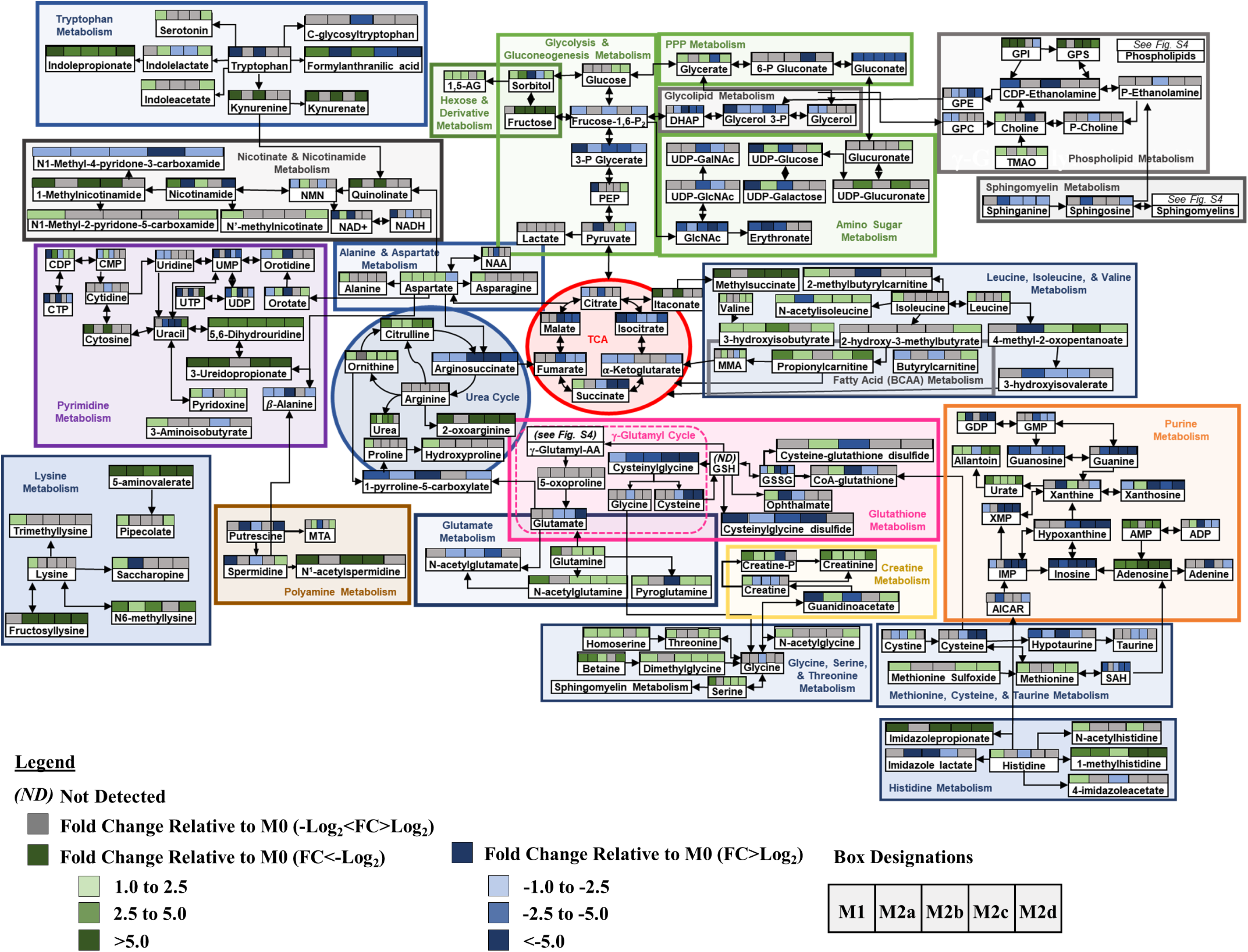
Schematic Demonstrating Metabolic Pathway Flux Relative to the Parent, Resting MΦ Phenotype (M0). Metabolic flux of selected metabolites within the specified pathways are denoted by arrow direction. Each of the colored boxes above the listed metabolite represents fold change relative to the parent M0 MΦs (1 < Log_2_ < -1). Fold change values (-Log_2_ <FC> Log_2_) are denoted in gray. Fold change values (-Log_2_ < FC) are denoted in shades of green and fold change values (FC > Log_2_) are denoted in gray shades of blue.

As MΦs are well characterized to be glycolytic, it was not surprising that *FBP1*, the gene encoding the FBPI regulator of gluconeogenesis, was significantly down regulated in all the polarized MΦs, especially M1 and M2b MΦs (Fig. 8A). Additionally, the accumulation of glucose observed in the M2c and M2d MΦs (Fig. 9: Glycolysis & Gluconeogenesis) suggests phenotypic variation in the utilization of glucose in MΦ immunometabolism. The accumulation of glycerate (Fig. 9: PPP) in the M1, M2b and M2d MΦs is suggestive of flux into the PPP for each of these phenotypes. From the topography analysis, the most statistically significant pathway observed was pyrimidine metabolism for both M1 and M2b MΦs (Fig. 7A & C; Fig. 9) while the purine pathway was most prevalent for M2d (Fig. 7E; Fig. 9). Furthermore, the upregulation of thioredoxin (encoded by *TXN*), a redox mediator, in M1 MΦs and the *CD44* transcript in M1 and M2b MΦs (Fig. 8A and 8D), known to enhance glycolysis and metabolic flux into the PPP [49], reinforced this pathway for immunometabolism in M1 and M2b MΦs.

One established metabotype feature of M1 polarization is the decoupled TCA after citrate and succinate, marked by accumulation of itaconate and succinate, which was observed in the M1 polarized MΦs (Fig. 9). Moreover, the upregulation of genes encoding an itaconate regulator, *IRG1* (Fig. 8A), and succinate-stabilized *HIF-1*α transcript (Supplemental Table 1) was consistent with these findings. Interestingly, M2b MΦs also exhibited similar *IRG1* and *HIF-1*α upregulation (Fig. 8A) along with itaconate accumulation; however, these MΦs did not accumulate succinate, contrasting with M1 MΦs. The remaining M2 MΦ subtypes (M2a, M2c, and M2d) all demonstrated metabolic flux patterns consistent with an intact TCA (Fig. 7-9).

For amino acid metabolism, the Arg & Pro and Ala, Asp, & Glu metabolic pathways were similarly impacted by polarization (p < 0.05) for all the MΦ phenotypes. In M1 MΦs, both glutamine and aspartate have been shown to replenish TCA metabolites such as α-ketoglutarate and fumarate, presumably depleted as a result of a decoupled TCA [28], and steady-state levels of both fumarate and malate were observed in the M1 MΦ, as well as accumulation of glutamine to feed into the TCA (Fig. 9). While only M1 MΦs accumulated alanine and asparagine, aspartate was accumulated in all polarized phenotypes apart from M2d MΦs (Fig. 9). Aspartate accumulation can drive β-alanine metabolism, which was a highly impacted metabolic pathway for the M1, M2a, M2b, and M2d MΦs (Fig. 7A, B, C, and E) and was consumed within these phenotypes during polarization (Supplemental Table 2 and Fig. 9: Pyrimidine). Aspartate can also lead to quinolinate production, as observed for M1 and M2b MΦs (Fig. 9: Nicotinate & Nicotinamide). In conjunction with upregulation of *CD38*, *IDO1*, and *NAMPT* (Fig. 8A & B), consumption of tryptophan, and accumulation of kynurenine (Fig. 9), these results reaffirmed previously findings highlighting the importance of tryptophan and nicotinamide metabolism within the M1 and M2b MΦs. In all M2 MΦ subtypes, taurine metabolism was impacted to various degrees during polarization, while cysteine and methionine metabolism were impacted by polarization in M2b, M2c, and M2d MΦ (Fig. 7B - E). Cysteine, taurine, and hypotaurine were generally consumed for the M2 MΦ subtypes (Fig. 9), with cysteine potentially feeding directly into glutathione metabolism in response to oxidative stress (Fig. 9). Moreover, diminished levels of oxidized glutathione and γ-glutamylamino acids, produced as a part of the γ-glutamyl cycle, further indicate oxidative stress associated with polarization across phenotypes (Fig. 9 and Supplemental Fig. 4A). Finally, Gly, Ser, and Thr metabolism was only identified as significant within the M1 MΦs (Fig. 7A), supporting previous findings in LPS-stimulated MΦs within mice [50].

Metabotyping in all five polarized MΦ phenotypes identified arginine metabolism as both impacted and significant (Fig. 7A-E). While intracellular stores of arginine did not appear to change with polarization, significant changes to metabolites upstream and downstream of arginine did, including arginosuccinate which feeds into the TCA Cycle, 1-pyrroline-5-carboxylate which derives from the γ-glutamyl cycle, and ornithine and citrulline of the Arg-Cit Cycle (Fig. 9: Urea Cycle). Interestingly, despite extensive literature around the expression of arginase (*Arg1*) as a marker of alternatively activated M2 MΦs, in our model expression of this gene was not significantly upregulated relative to the parent M0 MΦ phenotype (Fig. 8B); however, accumulation of ornithine in M1, M2a, M2c and M2d MΦs and urea in M1, M2a, M2b, and M2c MΦs suggest this enzyme is activated upon polarization (Fig. 9). Downstream of 1-pyrroline-5-carboxylate, proline and hydroxyproline accumulated in M1 MΦs, suggesting that the remaining M2 MΦ subtypes may utilize these metabolites for collagen biosynthesis (Fig. 9) [51]. Finally, arginine can be diverted to synthesize creatine through guanidinoacetate, which is highly consumed in both the M1 and M2b MΦs upon polarization (Fig. 9) and has been shown to suppress IFN-γ responses while facilitating IL-4 polarization in MΦs [52].

Other pathways of interest impacted by polarization included pyrimidine and purine metabolism (Fig. 7A- E). In general, these metabolic pathways were highly consumed across all MΦ phenotypes, but especially notable in the M1 and M2b MΦ consumption of dioxy- and trioxy-pyrimidines (Fig. 9). An offshoot of pyrimidine metabolism through β-alanine, polyamine metabolism was also markedly impacted with consumption of spermidine and accumulation of N1-acetyl-spermidine in both M1 and M2b MΦs upon polarization (Fig. 9). Hexosamine metabolism displayed a similar pattern with regards to M1 and M2b MΦ consumption; however, all three other M2 MΦ subtypes displayed metabolite accumulation, specifically of UDP-glucose, UDP-galactose, and UDP-glucuronate (Fig. 9). Whether these metabolites drive tissue regeneration/repair or act as signaling molecules within each MΦ phenotype, remains to be determined.

Finally, lipid metabolism was impacted in all five polarized MΦs, especially glycerophospholipid and sphingolipid metabolism, as shown by significance of impact (Fig. 7A-E). *COX-2* (*PTGS2*) transcript, encoding the enzyme involved in conversion of arachidonic acid to the pro-inflammatory precursor prostaglandin H2, was upregulated in M1 and M2b MΦs, while the transcript *ALOX5*, associated with anti-inflammatory response, were downregulated in M1 MΦs (Fig. 8D). Of the other alternatively activated M2 MΦ phenotypes, M2a MΦs upregulated *ALOX15*, which facilities activation of known M2a MΦ regulators (*PPARG*, *MRC1*, and *CD36*) (Figure 8C, E, and Supplemental Table 1) [53, 54]. M1 MΦ metabotype is also distinguished from the M2 MΦ subtypes by FA metabolism, with transcripts *CD36*, *ENPP2*, *FASN*, and *LPL* all downregulated with polarization and upregulated in M2a, M2c, and M2d to various levels (Fig. 8C). Lipid metabolite profiles reflected this distinction for the M1 MΦ metabotype, with general patterns of lipid accumulation for M1 MΦs and consumption of lipids for the M2 MΦ metabotypes, with the exception of DHA known to have anti-inflammatory properties (Supplemental Fig. 4B-F) [55]. Interestingly, the most dominant MΦ phenotype with regard to lipid metabolism was the M2b MΦ within which the levels of metabolite consumption upon polarization generally dwarfed all other phenotypes and, in contrast to similar myeloid functionality profiles, was clearly different from the M1 MΦ phenotype (Supplemental Fig. 4B-F). What impact this difference in lipid metabolism has on biological systems is under active investigation.

### MΦ Effector Functions and Metabotype Dynamics Over Time

Notorious for their plasticity, MΦ are rarely profiled over longer periods of stimulus and the *ex vivo* MΦ polarization model presented herein provided an excellent platform to compare polarization impact at different timepoints. To evaluate temporal shifts in both MΦ effector function and metabotype, polarized MΦ were harvested at both 24- and 72-hours polarization. Within the inflammatory protein subgroup, TNFα, CCL3, and CXCL8 exhibited minor significant increase only in TNFα from M1 MΦs at 72 hours polarization (Fig. 10A). While M1 MΦs markedly increased IL-12p70 and decreased IP-10 production between the two time points, IFNα production was decreased for M1, M2b, and M2c MΦ phenotypes (Fig. 10A). Among the inflammation regulating/tissue repair proteins (Fig. 10B) and growth factors (Fig. 10C), IL-10 and IL-6 production was significantly decreased from 24 to 72 hours polarization for both the M1 and M2b MΦ phenotypes. Interestingly, IL-1β concentrations increased dramatically for both phenotypes, while TGF-β concentrations decreased for all polarized MΦ phenotypes (Fig. 10B).

**Figure 10:**
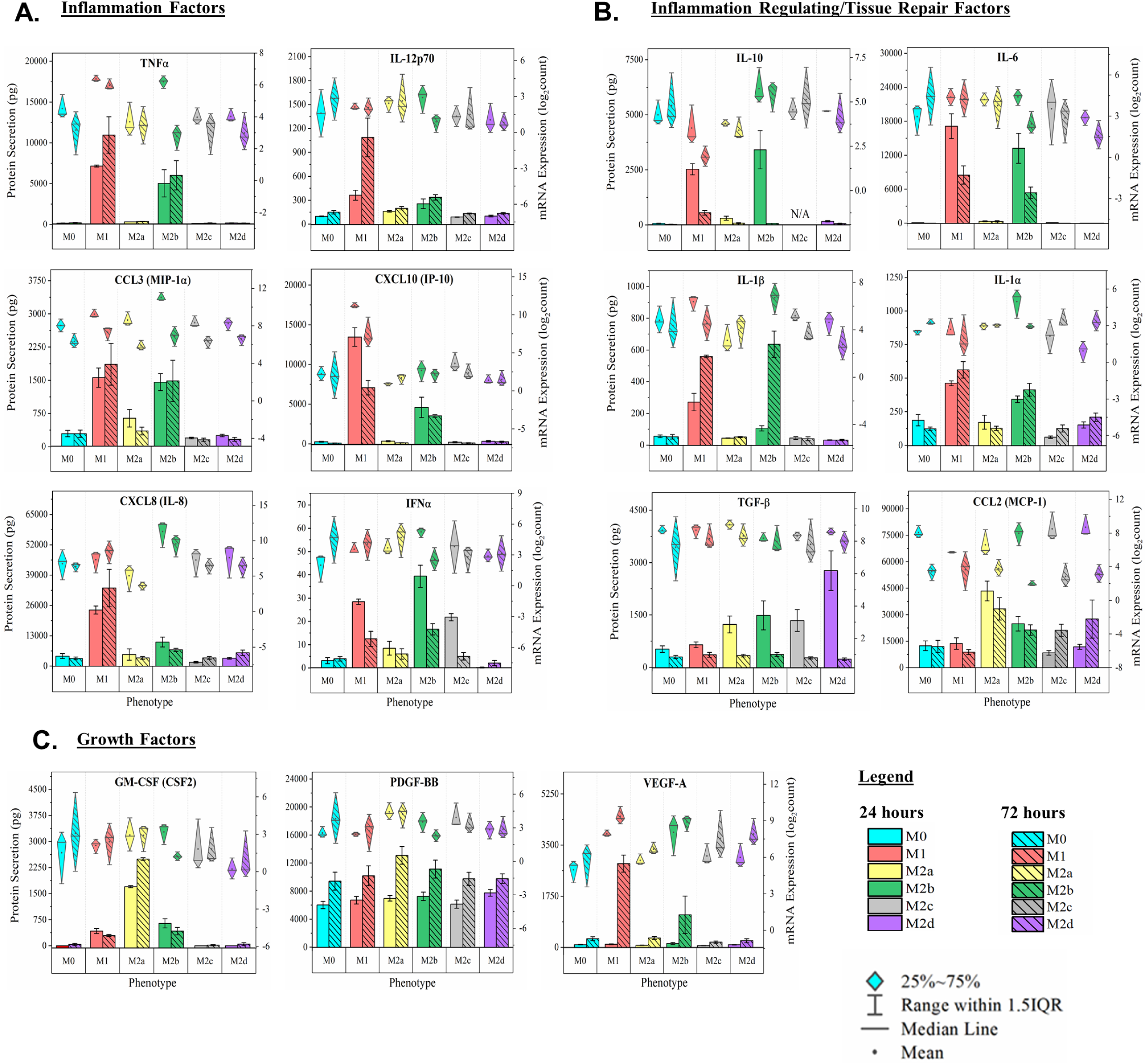
MΦ Functional Phenotyping at 72 Hours Post Polarization by Gene and Protein Expression of Immunomodulatory Factors. Multiplex detection of immunomodulatory factors in six MΦ phenotypes harvested at 72 hours were detected using magnetic bead-based quantification of mRNA and secreted protein. Immunomodulatory function includes pro-inflammatory (TNF⍺, IL-12p70, CCL3, CXCL10, CXCL8, and IFN⍺) **(A)**, immune-regulatory/tissue-repair (IL-10, IL-6, IL-1b, IL-1⍺, TGF-b, and CCL2) **(B)**, and growth factors (GM-CSF, PDGF-BB, and VEGF-A) **(C)** for the M0 resting MΦs (shown in blue), M1 MΦs (IFN-γ/LPS treated, shown in red), M2a MΦs (IL-4/IL-13 treated, shown in yellow), M2b MΦs (IC/LPS treated, shown in green), M2c MΦs (IL-10 treated, shown in gray), and M2d MΦs (IL-6/LIF treated, shown in purple). Diamond-whisker plots display 25%-75% quartile range, median, and mean. Bar charts indicate mean ± SEM. Expression profiles were normalized to total cell count and include three biological replicates (N=3).

Additionally, GO analysis demonstrated differentially expressed biological pathways between the 24- and 72-hour time points for all polarized phenotypes. M1 MΦs displayed a strong inflammatory signature at 24 hours, which was sustained at 72 hours (Fig. 11A, light red bars); however, by 72 hours, the M1 MΦs had initiated cellular protective/death processes such as the prostaglandin-endoperoxide synthase, negative regulation of apoptosis, and programmed necrotic cell death GO processes (Fig. 11A, dark red bars). For the M2a MΦs, GO processes at 24 hours heavily favored cellular proliferation, ECM remodeling, and angiogenesis (Fig. 11B, light yellow bars). While these cells still exhibited a potential for angiogenesis and ECM remodeling at 72 hours, the upregulation of the cyclooxygenase, leukotriene, histamine, and IFN-α processes at 72 hours were consistent with activation of anti-inflammatory function, viral inhibition, and the pathogenesis of asthma (Fig. 11B, dark yellow bar) [56]. As discussed above, M2b MΦs share a strong inflammatory functional signature with M1 MΦs at 24 hours (Fig. 11C, light green bars); however, by 72 hours, the M2b MΦs had upregulated genetic programs primarily associated with metabolism (Fig. 11C, dark green bars). ECM remodeling, angiogenesis, and efferocytosis functionality associated with M2c MΦs remained relatively consistent across both time points, with the interesting exception of IL-23 and IL-18 regulation, which are both generally associated with pro-inflammatory functionality. Finally, M2d MΦ functionality at 72 hours persisted with the angiogenic, profibrotic, and anti-inflammatory potential (Fig. 11D, light and dark purple bars); however, lipid metabolism was uniquely profiled at 72 hours, raising the interesting question of how this might translate to immunomodulation in this M2 MΦ subtype.

**Figure 11:**
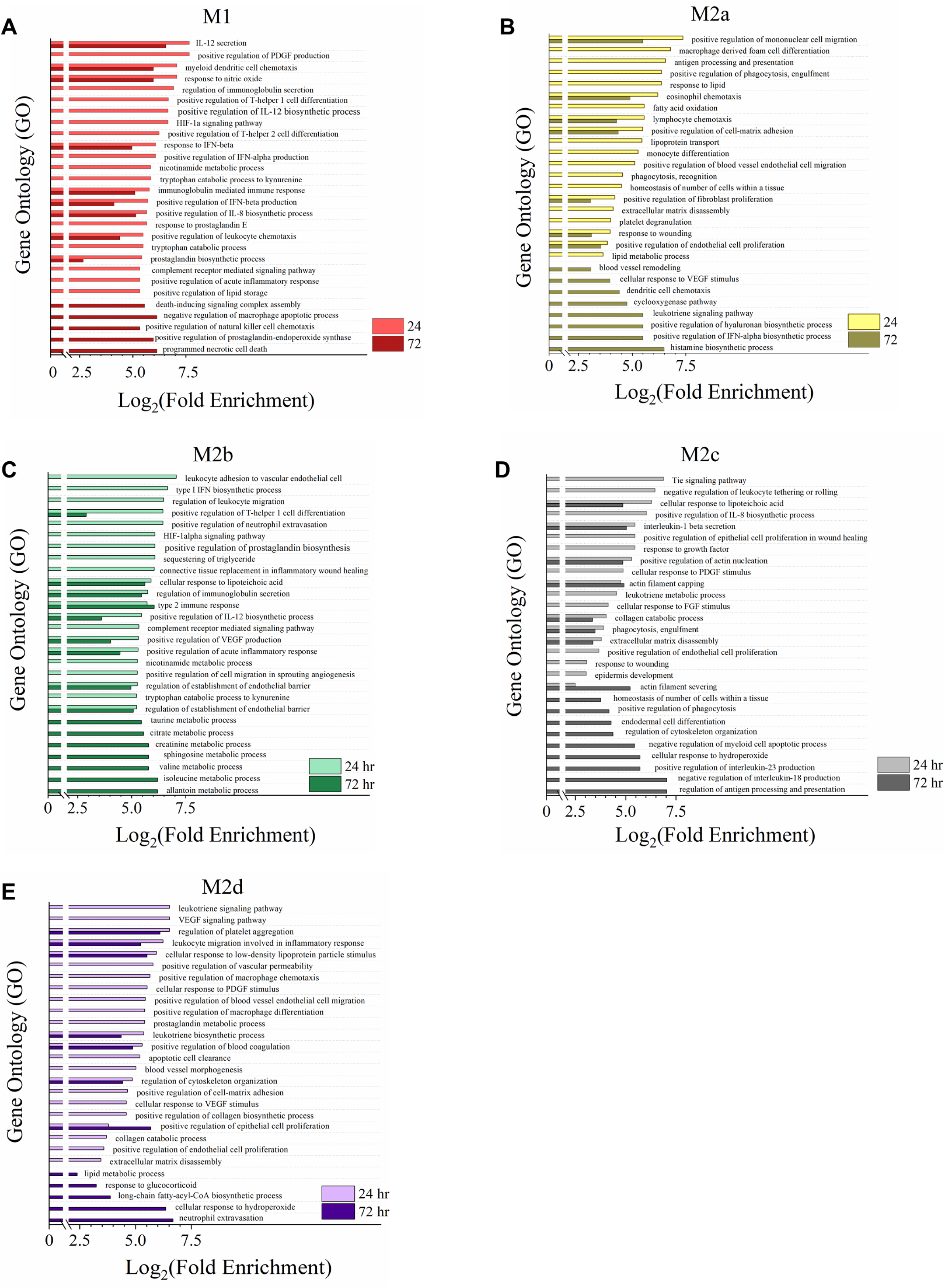
Gene Expression Profile Comparison of Polarized MΦ Phenotypes Identifies Unique and Overlapping Immunomodulatory Functions between 24- and 72-hour Polarization Times. Gene Ontology (GO) annotation of global myeloid gene expression identifies the differential biological functionality between the 24 (light colored bars) and 72-hour (dark colored bars) time points in five MΦ phenotypes. For each MΦ phenotype, the top 15 impacted GO processes significantly enriched (p ≤ 0.05) for both time points were plotted as such: M1 MΦs (IFN-γ/LPS treated, shown in red) **(A)**, M2a MΦs (IL-4/IL-13 treated, shown in yellow) **(B)**, M2b MΦs (IC/LPS treated, shown in green) **(C)**, M2c MΦs (IL-10 treated, shown in gray) **(D)**, and M2d MΦs (IL-6/LIF treated, shown in purple) **(E)**. GO process annotation was based on three experimental replicates for each MΦ s phenotype (N=3).

As described above, the top 15 impacted metabolic pathway scores were utilized to generate radar plots comparing 24- and 72-hour metabotypes in all five polarized MΦs (Fig. 12A-E, hatched and solid graphs for 24 and 72 hours, respectively). All phenotypes demonstrated significant shifts in metabolism from 24- to 72-hours polarization. For M1, M2a, M2b, and M2c MΦs, glycolysis and the PPP remained a significant part of the metabotype profile at 72 hours; however, glycolysis was significantly increased in the M2a MΦs (Fig. 11B), while remaining consistent in all of the other phenotypes. PPP metabolism also increased in the M1, M2a and M2b MΦs, but decreased in M2c MΦs. Additionally, amino acid and lipid metabolism was significant enhancement for both the M1 and M2b MΦs at 72 hours. The global differences in metabolic impact between the M2a (Fig. 11B) and M2c (Fig. 11D) MΦs is quite interesting in that they demonstrated nearly opposite metabotype profiles for the two time points. For example, M2a MΦs exhibited higher impact scores for several amino acids, including BCAA, as well as sphingolipid, galactose, glutathione, and purine metabolism at 24 hours (Fig. 11B, hatched plot), whereas these same pathways were more highly impacted at 72 hours for M2c MΦs (Fig. 11D, solid plot). Another interesting comparison is the different shifts in galactose, glutathione, purine, and pyrimidine metabolism in the M1 and M2b MΦs (Fig. 11A & C), further supporting the distinct metabotype profiles of these phenotypes despite their similar inflammatory signatures. Finally, in contrast to M1, M2a, and M2b MΦ metabolic dynamics, from 24 hrs. to 72 hours., M2d MΦs undergo a significant constriction in metabolic activity (Fig. 11E). This is especially evident when compared to the M2b MΦs, which significantly expand metabolic activity at 72 hours (Fig. 11C). While these observations in the dynamics of MΦ functionality and metabotype over time demonstrate that each of these MΦ phenotypes can exhibit significant plasticity, the causal relationship between metabolism and biological function remains an expansive area for future investigations.

**Figure 12:**
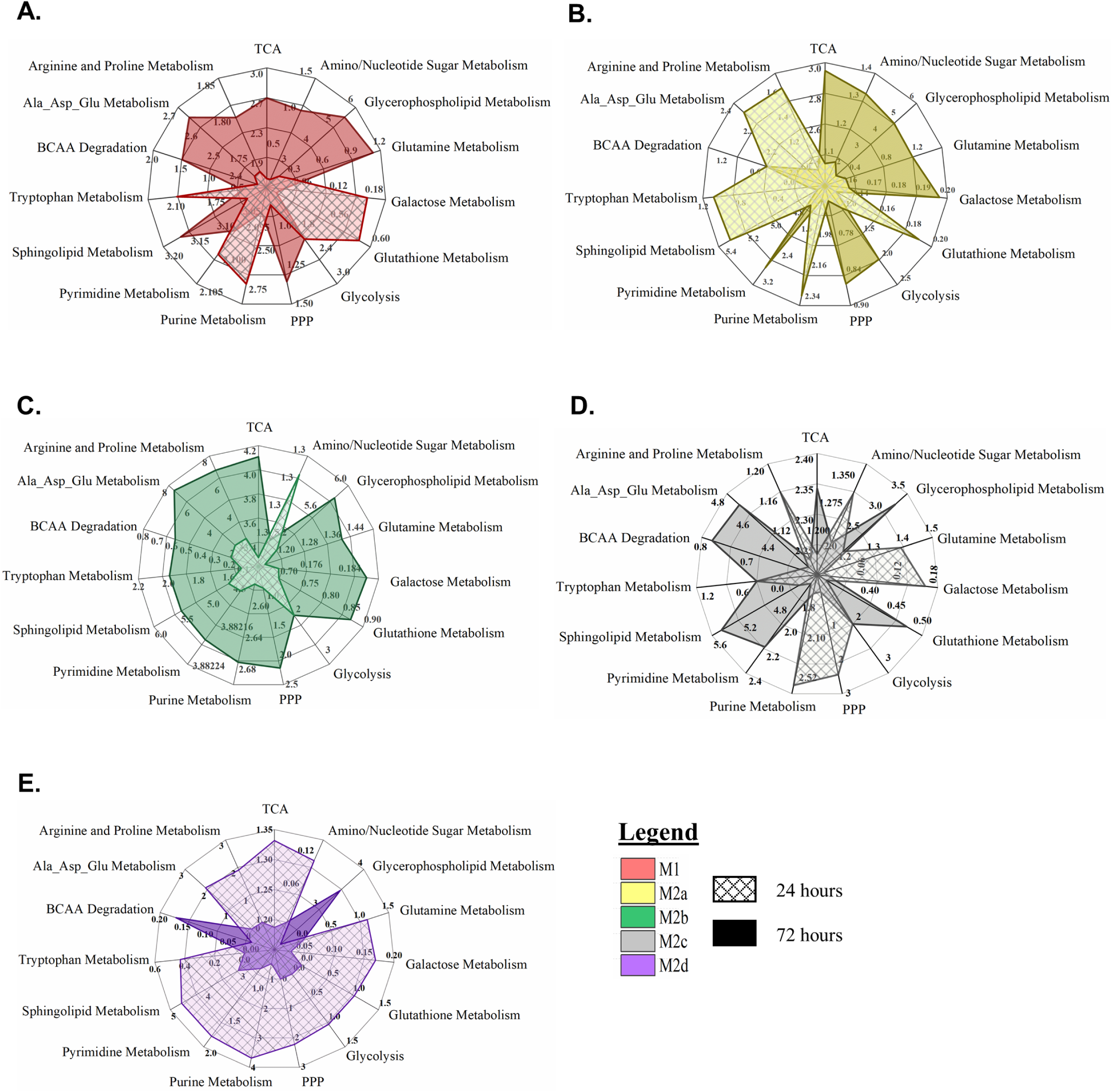
Metabotype Profiling of Polarized MΦs at 24 and 72 Hours. Pathway impact scores derived from pathway topology analysis were compared at 24 and 72 hours for the M1 MΦs (IFN-γ/LPS treated, shown in red) **(A)**, M2a MΦs (IL-4/IL-13 treated, shown in yellow) **(B)**, M2b MΦs (IC/LPS treated, shown in green) **(C)**, M2c MΦs (IL-10 treated, shown in gray) **(D)**, and M2d MΦs (IL-6/LIF treated, shown in purple) **(E)**. The top 15 enriched pathways are depicted in each spider plot with the hatched plots and solid plots representing the 24- and 72-hour time points, respectively.

## Discussion

Previous studies have described multiple MΦ phenotypes; however, to our knowledge, this study represents the most comprehensive profile and complete integration of metabotype within the known functional phenotypes. MΦ plasticity is uniquely dynamic and the integration of metabotype and functional phenotype presented herein provides a framework for further investigation into how this dynamic plasticity is driven. Furthermore, understanding how global metabolic shifts contribute to MΦ plasticity provides for greater insight into mechanisms of wound healing and provides the foundation for novel innovations in therapeutic design.

Exposure of resting M0 MΦs to IFNγ and LPS resulted in the classically defined M1 MΦ phenotype characterized by a strong, proinflammatory signature as evidenced by the secretion of several inflammatory cytokines and chemokines (Fig. 2A), the upregulation of M1-associated transcripts (Supplemental Table 1), and the biological functional profile identified through GO process analysis (Fig. 4). Furthermore, distinct metabolic shifts essential to this M1 MΦ functionality were also identified. Analysis of our metabolic data suggest reprogramming from OXPHOS to aerobic glycolysis, necessary for rapid ATP production and the provision of several glycolytic intermediates such as PKM2, GAPDH, and enolase for IL-1α/β, IL-6, and TNFα production (Fig. 2 & 9) [31, 57–59]. The diversion of glycolytic intermediates into the PPP drives the production of nucleotides, secondary metabolic intermediates, and the nucleotide substrate, NADPH [10, 28].

In parallel, pyruvate from glycolysis feeds into the observed decoupled TCA cycle. The first break before isocitrate results in an accumulation of citrate, which is diverted for itaconate production and *de novo* FAS. Upon LPS stimulation, the upregulation of *IRG1* (Fig. 8 and Supplemental Table 1) encodes the enzyme necessary for itaconate production (Fig. 5A, 9 and Supplemental Table 2) [60]. In addition to antimicrobial activity [61], itaconate serves as an inhibitor of SDH resulting in the accumulation of succinate (Fig. 9 and Supplemental Table 2) and the second break in the TCA cycle by M1 MΦs [60, 61]. The role of succinate as an inflammatory signal in MΦs is complex. Succinate oxidation through SDH together with an increased mitochondrial potential result in an increase of mitochondrial ROS production. Furthermore, accumulated succinate together with succinate recycling through GPR91 generates a feedforward loop that enhances HIF-1α stabilization (Supplemental Table 1) and the production of IL-1β (Fig. 2B) [27, 61, 62]. Additionally, excess citrate, once exported into the cytosol, can be converted into acetyl-CoA, oxidized to form malonyl-CoA, and converted into nascent FA products through the NADPH-dependent FASN [10]. Within the context of M1 polarization, the observed lipid accumulation (Supplemental Fig. 4) is essential for the synthesis of inflammatory mediators [63], prostaglandin synthesis through arachidonic metabolism (Fig. 8D) [64], and the membrane biosynthesis necessary for the phagocytic potential of these cells (Fig. 1) [65].

For the M2a MΦs, our metabolic results support both active glycolysis and an intact TCA cycle (Fig. 9 and Supplemental Table 2) although glucose does not appear to be consumed at the same rate as it was in the M1 phenotype. While M2a MΦs demonstrate active glycolysis, which is used to supply the TCA cycle, M2a polarization has been shown to be unaffected during glucose deprivation. This observation suggests that other metabolic pathways, such as glutaminolysis, can also be utilized to support the TCA cycle and drive oxidative metabolism [65]. We observed decreased pools of lipids in this MΦ subtype (Supplemental Fig. 4), suggesting that FAO is indeed active. Previous research has shown that glucose oxidation and mitochondrial respiration are necessary for early IL-4 polarization, but not FAO; however, FAO, dependent on CD36 expression (Fig. 8C) [32], was established as critical for M2a functionality at later time points [33]. Furthermore, suppression of CD36 FA uptake along with FAO has been shown to ameliorate M2a polarization during parasitic helminth infections [32]. Using metabolic flux analysis and glutamine deprivation, Jha, et al., (2015) demonstrated the necessity of glutamine for amino and nucleotide sugar metabolism and M2a polarization markers CD301 and MRC1 [28]. While glutamine flux analysis is outside of the scope of the research presented herein, we did observe increased metabolic activity within the hexosamine biosynthesis pathway and increased pools of UDP-GlcNAc-associated modules (Fig. 9 and Supplemental Table 2).

Much of what is known of the biologically functionality of the M2a phenotype lies within the context of wound healing. In the late stages of inflammation, M2a MΦs facilitate the clearance of glycosylated pathogens through the upregulated cell-surface MRC1 (Supplemental Table 1) and the efferocytosis of apoptotic cells in a CD36-dependent fashion [66]. Additionally, MRC1 is known to target mannosylated antigen intracellularly for MHC-II compartments and antigen presentation in dendritic cells [67]. Interestingly, our M2a polarized MΦs show increased expression of the MHC-II receptor HLA-DR (Fig. 1D & H) suggesting a similar role for this MΦ subtype. In addition to pathogen clearance, the MRC1 removes glycosylated inflammatory proteins from the wound microenvironment [68]. M2a MΦs also upregulate and secrete numerous biological factors, such as IL-1RA (Supplemental Fig. 1G) to suppress IL-1 mediated inflammation [17], the chemokine ligands CCL3 (Fig. 2A), CCL17, CCL18, and CCL22 (Supplemental Table 1), and growth factors GM-CSF, TGF-β, and PDGF (Fig. 2B & C) needed to promote the recruitment and differentiation of numerous cell types necessary for the proliferative phase of wound healing [69].

Within a simplistic MΦ polarization paradigm, M1 MΦs promote inflammatory responses while M2 MΦ- associated phenotypes drive anti-inflammatory, wound healing functions; however, resting MΦs exposed to TLR-agonist, endogenous ligands coupled with FCγ receptor ligation and signaling through SYK resulted in a polarized phenotype that defies the M1/M2 paradigm by secreting strong pro-inflammatory cytokines while promoting biological effector mechanisms associated with the M2 family of MΦs [20, 70]. Indeed, Vogelpoel, et al. (2014) demonstrated that TLR agonists alone, using both LPS and Pam3CSK4, could not produce this phenotype that is commonly known as M2b MΦs [8]. Using polarization stimulus through LPS and IC, this MΦ phenotype was demonstrated to secrete proinflammatory cytokines TNFα, IL-1β, and IL-6 (Fig. 2A & B) and upregulated transcripts, including *TNF*, *IL-1β*, *IL-23A*, *IL-10*, *CCL1*, and *CCL5* (Supplemental Table 1), that promote Th17 responses [9] and profile as distinct from any other M2 MΦ subtype.

Metabolically, M2b MΦs displayed similar glycolysis, tryptophan, amino sugar, and pyrimidine metabolism as M1 MΦs; however, in contrast to M1 MΦs, the TCA cycle appears to be intact in M2b MΦs, as shown by the lack of succinate accumulation (Fig. 9). In LPS-activated MΦ, HIF-1α stabilization and IL-1β production has been linked to succinate accumulation via glutamine-dependent anaplerosis or the γ-aminobutyric acid (GABA) shunt [27]; however, this does not appear to be an active mechanism in M2b MΦs and thus warrants further investigation. Additionally, the M2b MΦ lipid metabolite profile suggests active FAO or OXPHOS similar to the other M2 subtypes (Supplemental Fig. 4). M2b MΦs demonstrate complex functionality. For example, the inflammatory signature can be readily seen contributing to the pathology of chronic inflammatory conditions such as rheumatoid arthritis [8] and SLE [9]; however, M2b MΦs have also been associated with aberrant antimicrobial responses [21, 71], reduced cardiac fibrosis after ischemia/reperfusion injuries [72], and improvement of intestinal colitis [73].

IL-10, the polarizing stimuli for M2c MΦs in this study, is a known inducer of CD163 expression that has historically defined this functional phenotype [23]; therefore, it was not surprising that cell-surface marker analysis characterized this M2 MΦ subtype as CD86^low^CD163^high^ (Fig. 1). These findings, along with the observed upregulation of *MerTk* (Supplemental Table 1), a key component of efferocytosis, is consistent with phagocytosis GO processes being profiled (Fig. 4). CD163, the hemoglobin scavenger receptor, mediates Hb-oxidative tissue damage following hemolysis and induces the expression of the anti-inflammatory enzyme heme oxidase 1 (*HMOX1;* data not shown) [74]. Additionally, CD163 expression accompanied by other anti-inflammatory mediators was observed in cardiac bypass patients post-surgery and in volunteers recovering from cantharidin-induced skin blisters [75]. Our findings are in line with these studies and are consistent with a functional phenotype responsible for the regulation and mediation of inflammation, vascular insult, and oxidative stress [15, 74, 75].

To date, extraordinarily little is known regarding the metabolism of M2c MΦs and has been limited to carbohydrate and glutamate metabolism. Rodriguez-Prados, et al. (2010) concluded that M1, M2a, and M2c polarized MΦs were glycolytic cells that converted up to 95% of glucose into lactate. While this conversion was accelerated for the M1 cells, both M2a and M2c MΦs exhibited basal glucose consumption levels comparable to the M0 parent cells [30]. Interestingly, our results demonstrated that M2c MΦs accumulated a significant amount of glucose relative to the M0 MΦs (Fig. 9 & Supplemental Table 2), indicating either a slower glycolytic capacity or the ability the import glucose at higher concentrations. Additionally, these authors found differences in glutamine consumption (M1>>M2a>>M0>M2c) and glutamate production (M1>M2c>M2a>M0) in their study [30]. While we observed different accumulations of glutamine and glutamate in these phenotypes, the M2c MΦs displayed considerably diminished pools of glutamate in comparison to the M0 parent cells (Fig. 9 & Supplemental Table 2). Given the necessity of glutamate in ECM remodeling, it is probable that glutamate is being diverted to collagen synthesis (as reviewed in Karna, et al. 2020 [76]). In addition to active glycolysis, an observed shift to the PPP via glucuronate and the accumulation of UDP-glucuronate (Fig. 9 and Supplemental Table 2) suggests a potential investment in the synthesis of hyaluronic acid for tissue regeneration and anti-inflammatory responses during wound healing [77]. M2c MΦs also demonstrated an intact TCA cycle, enhanced OXPHOS, and lipid catabolism (data not shown and Supplemental Fig. 4). There also appears to be concurrent urea and aspartate-arginosuccinate cycles (Fig. 9) as well as investment in BCAA metabolism (Fig. 9).

Given the remarkable plasticity of MΦs and the equally disparate array of tumor-specific microenvironments, it is not surprising that TAMs, as a collective, exhibit diverse functionality resembling both M1 and M2 polarized phenotypes [78]. As such, TAMs have been associated with protein expression patterns observed in both acute wounding (i.e. IL-6, and IL-1β) and wound resolution (i.e. TGF-β, PDGF and VEGFA) [26]. IL-6 appears to be central to many of the underlying processes that drive TAM functionality. Muliaditan, et al. (2018) demonstrated that TAM-produced IL-6 regulates the expression of *HMOX1* which has been associated with poor overall survival rates. In our IL-6 polarized M2d MΦs, we also observe the upregulation of *HMOX1* (data not shown), supporting these findings [26]. In terms of overall biological functionality, TAMs have been shown to facilitate angiogenesis [79], tumor progression [6], immunosuppression [25], and therapeutic resistance [5]. Our transcriptional profile of these MΦs directly supports all these functions (Fig. 4). Furthermore, TGF-β and PDGF-BB (Fig. 2C), secreted at significant levels in these cells, are well-documented factors associated with tumor progression and generally poor outcomes [80].

The tumor metabolic environment typically has some degree of hypoxia, which promotes TAM functional plasticity in support of tumor survival. This hypoxia shifts TAMs towards oxidative metabolism and promotion of neoangiogenesis and metastasis [79]. In our model, M2d MΦs were not cultured under hypoxic conditions and glucose accumulation was observed (Fig. 9), indicating functional glycolysis, as has been observed in primary tumors [4, 81, 82]. The tumor environment has also been characterized as abundant in lipids and M2d MΦs were observed to highly express CD36, the fatty acid translocase integral membrane protein (Fig. 7). Interestingly, in a murine melanoma model, CD36 expression on TAMs was found to mediate uptake of oxidized LDL and promote tumorigenesis [83]. FAO utilization in TAMS has also been proposed to be mediated through epigenetic reprogramming and pharmacological inhibition of FAO favors M2-to-M1 repolarization in murine cancer models [25, 84]. In our model system, the M2d MΦs displayed significant FAO, indicating a key role for this functional phenotype (Fig S8); however, investigation into potential epigenetic modifications cellular cross-talk with tumorigenic cells is outside of the scope of the present study.

Finally, macrophage functional plasticity and immunomodulatory effects of metabolism have rarely been observed over an extended period. By extending our *ex vivo* MΦ polarization model out to 72 hours post polarization, the temporal effects of stimulus were observed. While pro-inflammatory MΦs (M1) shifted to programmed cell death and anti-inflammatory MΦs (M2a and M2c) continued to promote tissue remodeling/repair function as would be expected with extended stimulus, M2b and M2d MΦs demonstrated significant and opposite shifts in metabolic programming. For example, apart from glycerophospholipid metabolism and BCAA degradation, the M2d MΦs displayed a massive contraction of metabolic activity, while the M2b MΦs display a massive increase in metabolic activity (Fig. 12C, E). The functional consequences of this metabolic shift are suggested by down regulation of TGFβ in M2d MΦs and up regulation of IL-1β in M2b MΦs; however, further investigation is warranted and ongoing.

In conclusion, this study presents the most comprehensive functional phenotyping of MΦ polarization plasticity and associated metabotypes to date. While the *ex vivo* MΦ model system utilized primary, human blood derived MΦs and allowed homogeneous polarization of MΦ phenotypes, the functional phenotypes and associated metabotypes presented herein must be established *in vivo* and in functional context. *In situ* tissue profiling within primary, human wound tissue is ongoing; however, demonstrating the potential *in vivo* context across numerous tissues and pathologies provides unlimited opportunity for investigation. In addition, the work presented herein demonstrates correlation between MΦ functional phenotype and metabotype but deciphering the intricate interplay between metabolism and immunomodulation at the heart of causation is outside of the scope of the present investigation. Clearly, next steps must include contextualizing these profiles and teasing out the mechanistic structure of metabolic immunomodulation of MΦ functional plasticity.

## Authorship

C.B.A, M.C.B.A, and M.M.D. conceptualized and designed the experimental approach and supervised sample collection for this research. C.B.A, T.M.W.L., H.L.S., and J.G. contributed equally to the execution of the experiments and data collection. C.B.A and M.C.B.A. performed the data analysis, generated the figures, and wrote the manuscript. All authors participated in the editing process.

## Acknowledgments

This work was supported in part by the U.S. Department of Veterans Affairs, Office of Research and Development Biomedical Laboratory Research Program, the Idaho INBRE Program (NIH NIGMS P20 GM103408), a NIAID-NIH award (RO3AI135998; PI Ammons), and the IVREF Center of Biomedical Research Excellence in Emerging/Reemerging Infectious Diseases (NIH NIGMS P20GM109007). This content is solely the responsibility of the authors and does not necessarily represent the official views of the National Institutes of Health, U.S. Department of Veterans Affairs, or the United States Government.

## Abbreviations

αKG: alpha ketoglutarate
AHS: autologous human serum
ALA: alanine
AMP: adenosine monophosphate
ANOVA: analysis of variance
APC: allophycocyanin
APCs: antigen-presenting cells
ARG: arginine
ASP: aspartate
ATP: adenosine triphosphate
BCAA: branch chain amino acid
bFGF: basic fibroblast growth factor
BSA: bovine serum albumin
CCL: chemokine (C-C motif) ligand
CD: cluster of differentiation
CIT: citruline
CSV: comma-separated value
CXCL: chemokine (C-X-C motif) ligand
CYS: cysteine
DAMPs: damage-associated molecular patterns
DGS: directed global significance
ECM: extracellular matrix
EGF: epidermal growth factor
FAO: fatty acid oxidation
FAs: fatty acids
FASN: fatty acid synthase
FDR: false discovery rate
FGF: fibroblast growth factor
FITC: fluorescein isothiocyanate
FMO: fluorescence minus one
Fcγ: fragment crystallizable gamma region
GLU: glutamate
GLY: glycine
GM-CSF: granulocyte-macrophage colony-stimulating factor
GO: gene ontology
GSEA: gene set enrichment analysis
Hb: hemoglobin
HC: hierarchical clustering
HESI-II: heated electrospray ionization
HIF: hypoxia-inducible factor
HIS: histidine
HLA-DR: human leukocyte antigen – DR isotype
ICs: immune complexes
IFN-γ: interferon gamma
IL: interleukin
IL-1Ra: IL-1 receptor agonist
IRB: institutional review board
LIF: leukemia inhibitory factor
LPS: lipopolysaccharides
M-CSF: macrophage colony stimulating factor
MEM: minimum essential media
MerTK: Mer receptor tyrosine kinase
MFI: median fluorescence intensity
MHC-II: major histocompatibility complex class II
MMP: matrix metalloproteases
MRC1: mannose receptor
MΦs: macrophages
NADPH: nicotinamide adenine dinucleotide phosphate
NO: nitric oxide
OPLSDA: orthogonal PLSDA
OXPHOS: oxidative phosphorylation
PBMCs: primary blood-derived mononuclear cells
PBS: phosphate buffered saline
PCA: principal component analysis
PDGF: platelet-derived growth factor
PE: phycoerythrin
PerCP-Cy: peridinin chlorophyll protein complex cyanine
PFPA: perfluoropentanoic acid
PGF: placental growth factor
PLSDA: partial least-squares to latent structures discriminant analysis
PPP: pentose phosphate pathway
PRO: proline
QC: quality control
RNS: reactive nitrogen species
ROS: reactive oxygen species
RPMI: Roswell Park Memorial Institute
RSD: relative standard deviation
SDEGs: significantly differentially expressed genes
SDH: succinate dehydrogenase
SER: serine
SLE: systemic lupus erythematosus
SYK: spleen tyrosine kinase
SPMs: specialized pro-resolving mediators
SUS: Shared and Unique Structure
T: helper cells Th cells
TAMs: tumor associated MΦs
TCA: tricarboxylic acid cycle
TGF-β: transforming growth factor-β
THR: threonine
TLR: toll-like receptor
TNF: tumor necrosis factor
TXN: thioredoxin
UDS: undirected global significance
UHPLC/MS/MS: ultra-high-performance liquid chromatography/tandem accurate mass spectrometry
VEGF: vascular endothelial growth factor

---

**Supplemental Figure 1:**
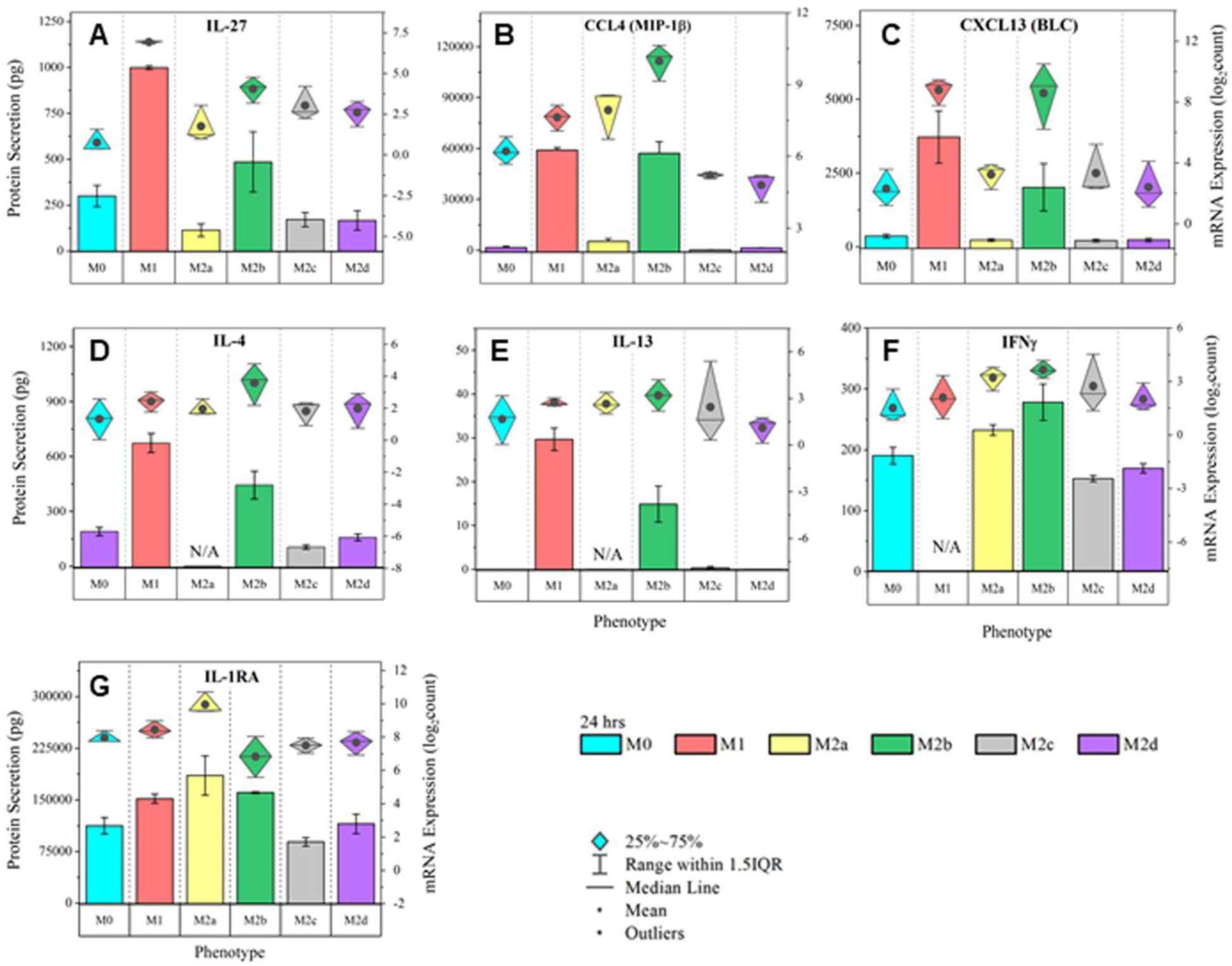
MΦ Functional Phenotyping by Gene and Protein Expression of Immunomodulatory Factors. Multiplex detection of immunomodulatory factors in six MΦ phenotypes were detected using magnetic bead-based quantification of mRNA and secreted protein. CD14+ monocytes were isolated from human blood-derived PBMCs, differentiated into resting MΦ (M0, shown in blue) with M-CSF *ex vivo*, and polarized into five activated phenotypes using IFN-γ/LPS (M1, shown in red), IL-4/IL-13 (M2a, shown in yellow), IC/LPS (M2b, shown in green), IL-10 (M2c, shown in gray), or IL-6/LIF (M2d, shown in purple). Gene and protein expression profile (diamond-whisker plots and bar charts, respectively) of key functional molecules were profiled by multiplex assay after 24 hours of *ex vivo* polarization. Immunomodulatory factors include IL-27 **(A)**, CCL4 **(B)**, CXCL13 **(C)** IL-4 **(D),** IL-13 **(E),** IFNγ **(F),** and IL-1RA **(G).** Diamond-whisker plots display 25%-75% quartile range, median, and mean. Bar charts indicate mean ± SEM. Expression profiles were normalized to total cell count and include three biological replicates (N=3).

**Supplemental Figure 2:**
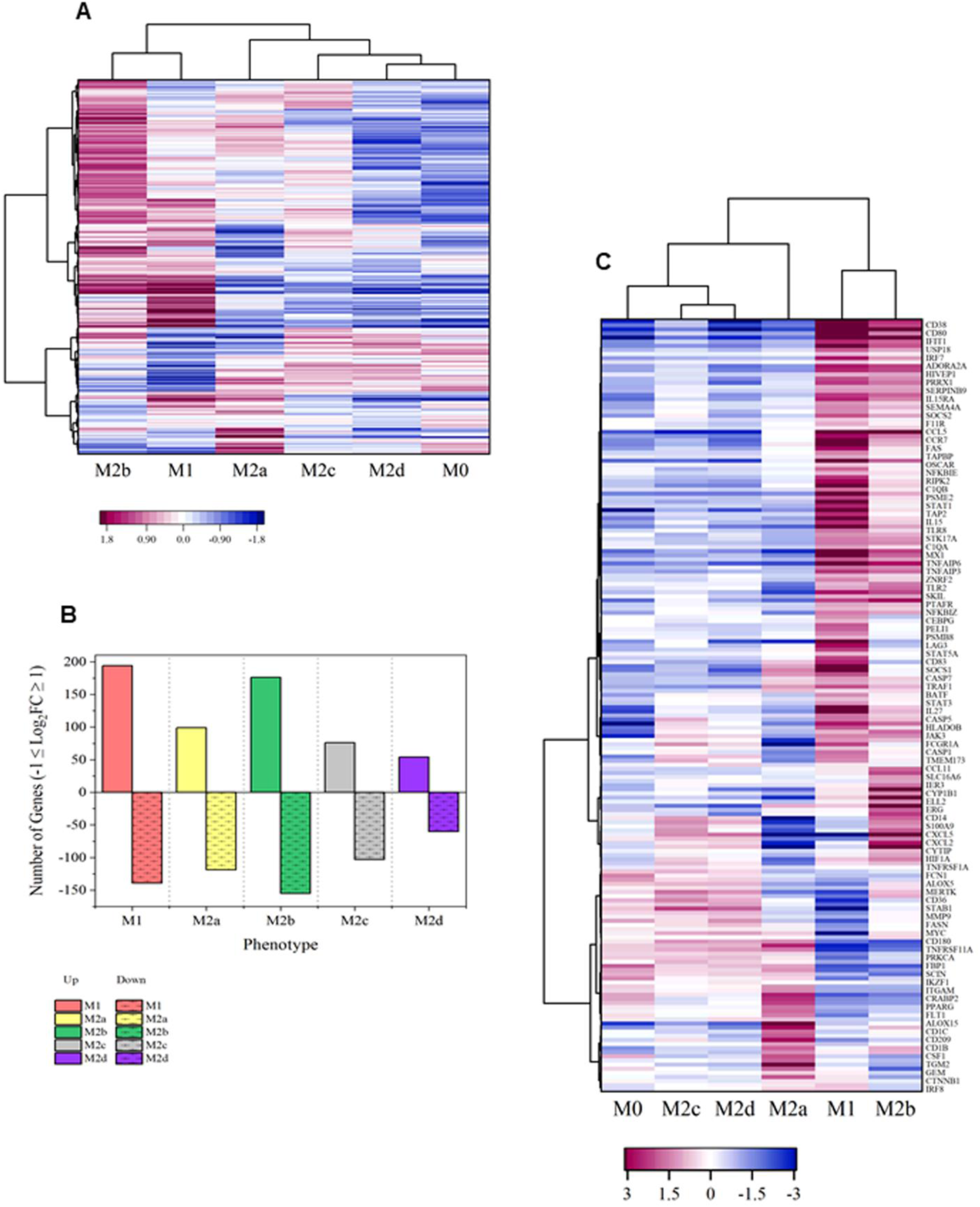
Global Transcriptomics Differentiates the MΦ Functional Phenotypes from the Parent, Resting M0 Phenotype. Hierarchical cluster analysis performed on the normalized expression data identified differences in the transcriptional profiles of the polarized MΦ phenotypes relative to the M0 parent MΦ. Distance measure is by Pearson correlation with clustering determined using the Ward algorithm. Heatmap scale (-1.8 to 1.8) is colored from blue to red **(A).** The fold change for each gene was calculated relative to the M0 phenotype and the number of up- and down-regulated genes (-1≤ Log_2_FC ≥1) graphed in **(B).** Gene set enrichment analysis (GSEA) of the global myeloid gene expression revealed 189 significantly differentially expressed genes (SDEGs) identified using a one-way ANOVA with an adjusted p-value (FDR) cutoff of 0.05. Hierarchical cluster analysis was performed on this data using a distance measure calculated using Pearson correlation with clustering determined using the Ward algorithm. Heatmap scale (-1.8 to 1.8) is colored from blue to red **(C).**

**Supplemental Figure 3:**
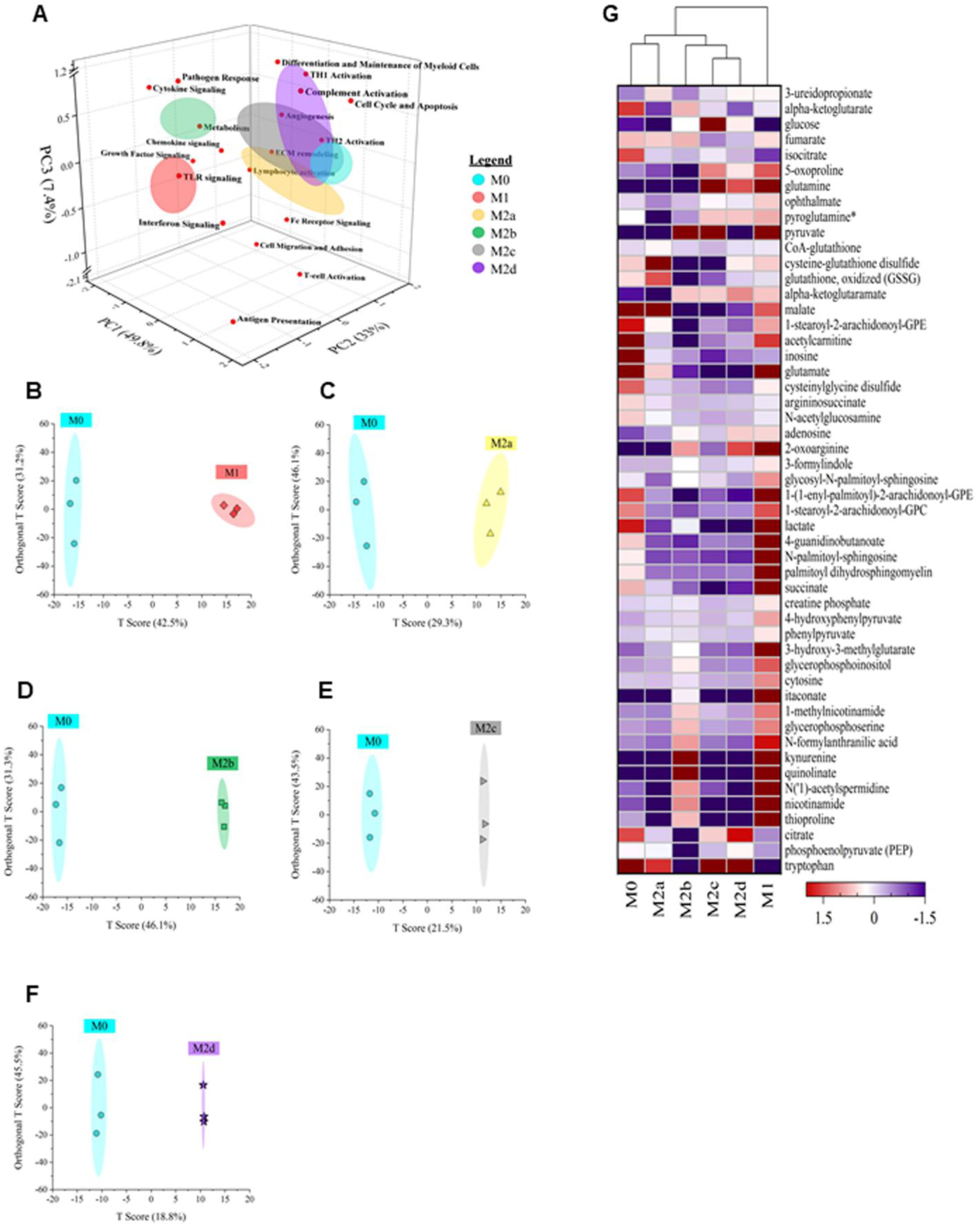
MΦ Polarization into Functional Phenotypes Activates Distinct Gene Expression Profiles and Metabolic Differences Relative to the Parent, Resting MΦ Phenotype (M0). Principal component analysis (PCA) 3D loadings plot utilizes red dots to depict the PC1 (x-axis), PC2 (y-axis) and PC3 (z-axis) loadings for each of the 19 Nanostring identified pathways while the colored ellipses denote the 3D representation of the linear combination of the three principal components for each polarized phenotype and the M0 parent cells **(A)**. Orthogonal Projections to Latent Structures Discriminant Analysis (OPLSDA) clustering scores plots with 2D T-scores are depicted for The M0 parent cells (cyan) versus M1 MΦs (IFN-γ/LPS treated shown in red) **(B)**, M2a MΦs (IL-4/IL-13 treated, shown in yellow) **(C)**, M2b MΦs (IC/LPS treated, shown in green) **(D)**, M2c MΦs (IL-10 treated, shown in gray) **(E)**, and M2d MΦs (IL-6/LIF treated, shown in purple) **(F).** Global metabolomics profile of the parent, resting MΦ phenotype (M0) and five polarized MΦ phenotypes identified 498 compounds of known identity, normalized to cell count. Hierarchical cluster analysis was performed on the top 50 significantly differentiated metabolites identified using a one-way ANOVA with an adjusted p-value (FDR) cutoff of 0.05 using a distance measure calculated using Pearson correlation with clustering determined using the Ward algorithm. Heatmap scale (- 1 to 1) is colored from purple to red **(G).**

**Supplemental Figure 4:**
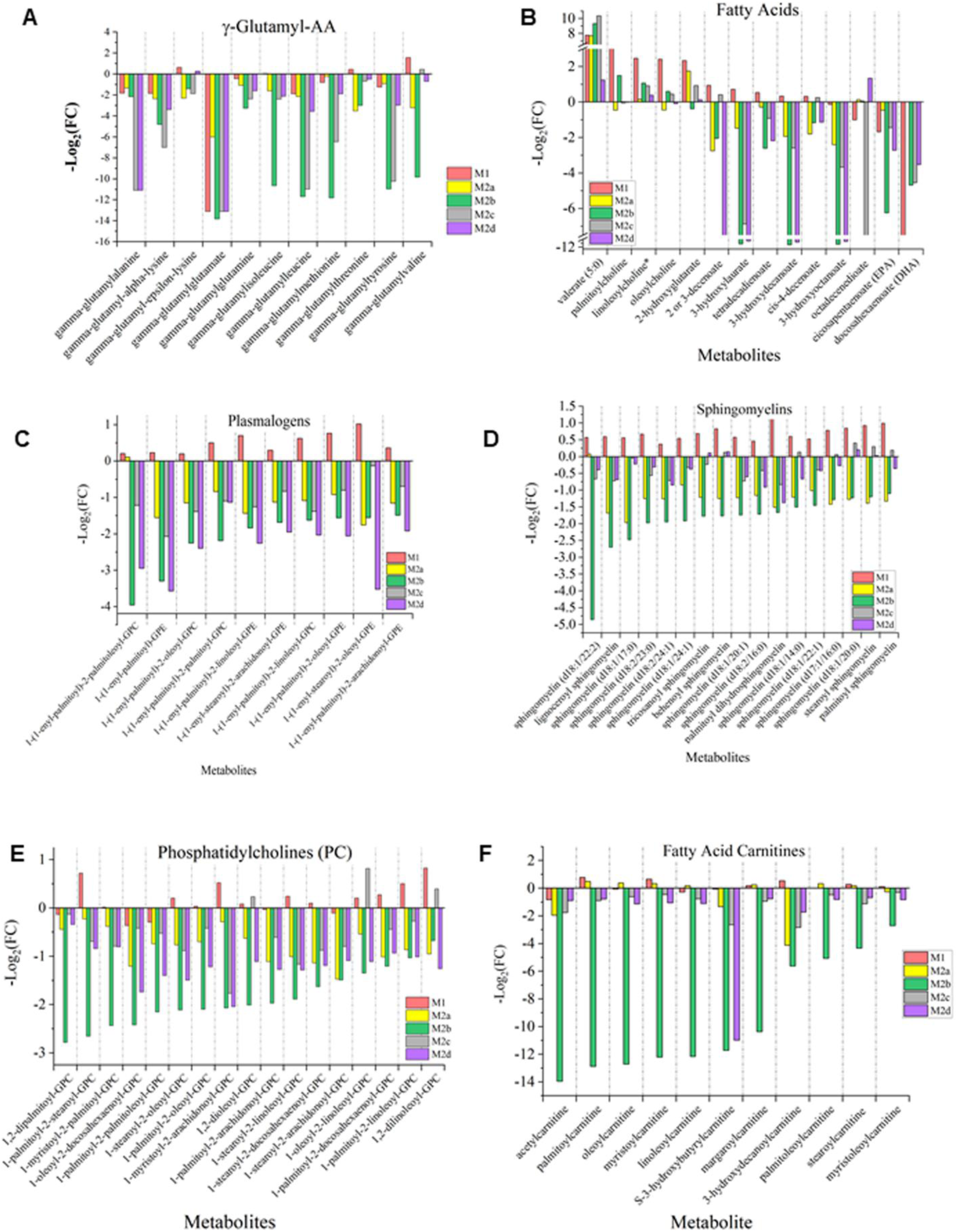
Untargeted Metabolomics of Five Polarized MΦ Phenotypes Identifies Phenotypic Differences in γ-Glutamyl Amino Acid Metabolism and Lipid Metabolism Relative to the Parent, Resting MΦ Phenotype. Histogram plots depict the accumulation of (positive –Log_2_FC bars) or decrease of (negative –Log_2_FC bars) lipid metabolites for the M1 MΦs (IFN-γ/LPS treated, shown in red), M2a MΦs (IL-4/IL-13 treated, shown in yellow), M2b MΦs (IC/LPS treated, shown in green), M2c MΦs (IL-10 treated, shown in gray), and M2d MΦs (IL-6/LIF treated, shown in purple) relative to the M0 parent cells. The represented metabolic classifications include γ-glutamyl amino acids **(A),** fatty acids **(B)**, plamologens **(C)**, sphingomyelins **(D)**, phosphotidylcholines (PC) **(E)**, and fatty acid carnitines **(F)**.

**Supplemental Table 1:**
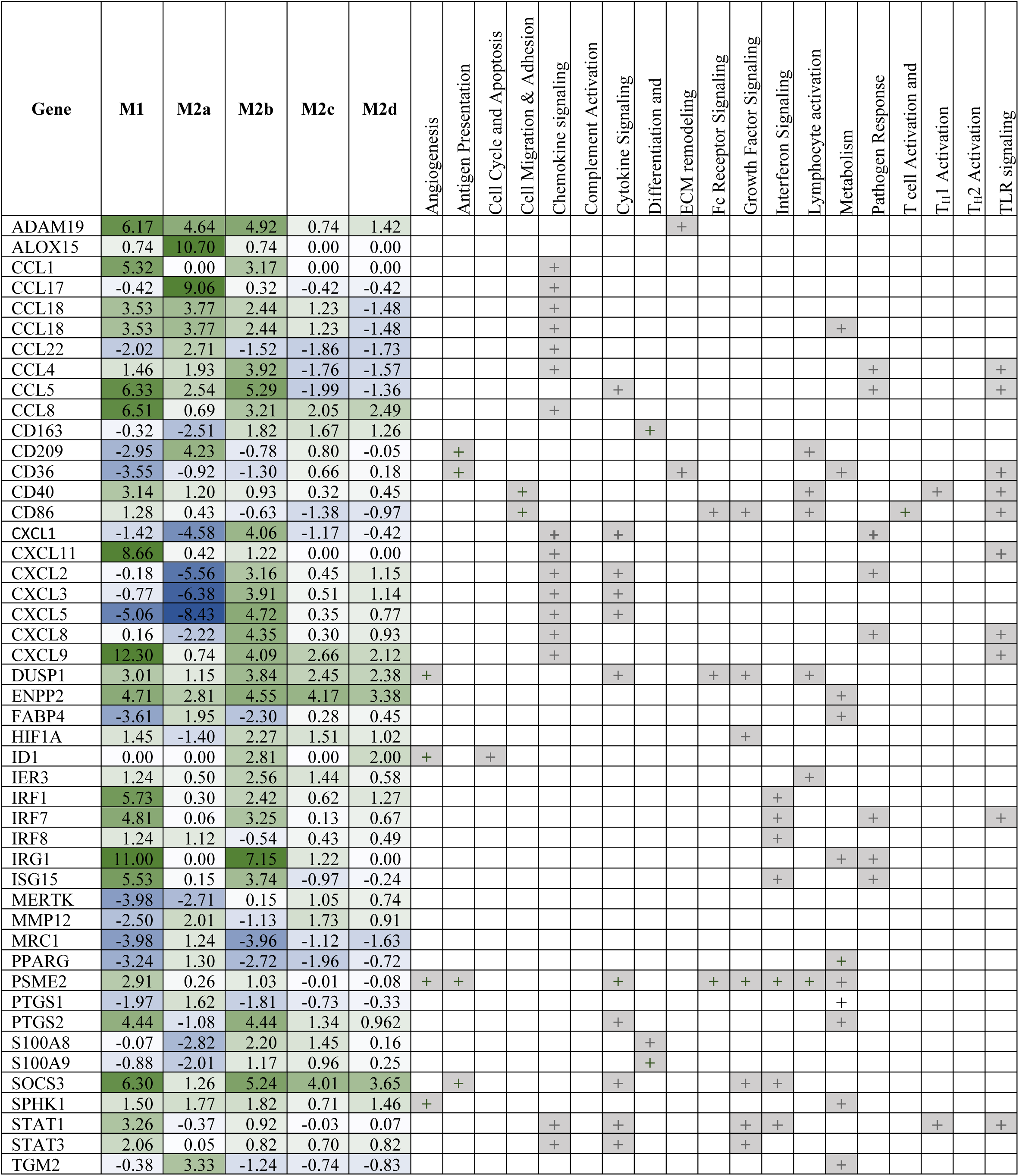
Relative genetic expression of select transcripts for each MΦ polarized phenotype compared to the Parent, Resting MΦ Phenotype (M0). Heatmap scale (-8 to 8) is colored from blue to green. Nanostring-defined pathways associated with specific SDEGs are denoted with gray box with a plus sign.

**Supplemental Table 2:**
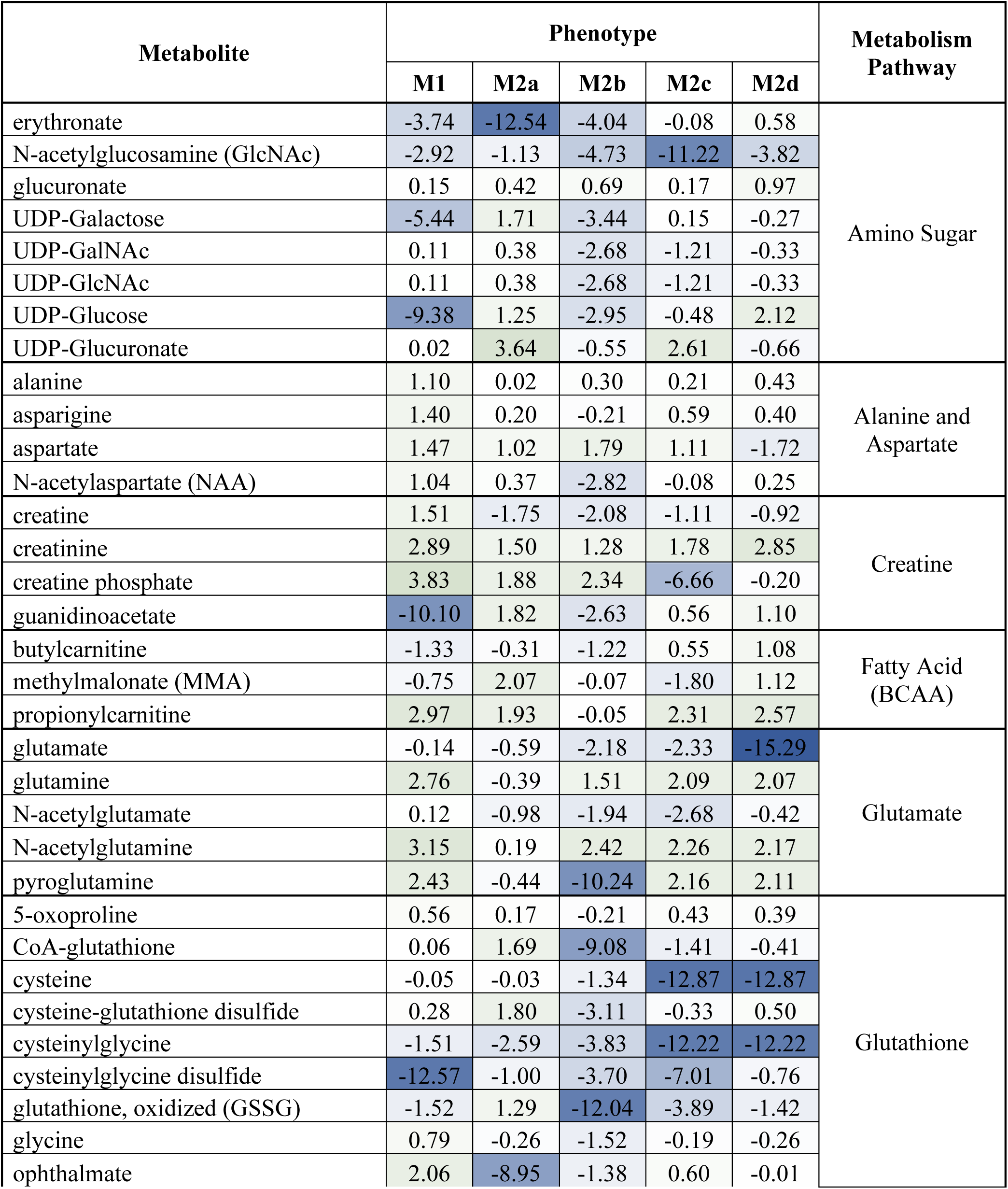
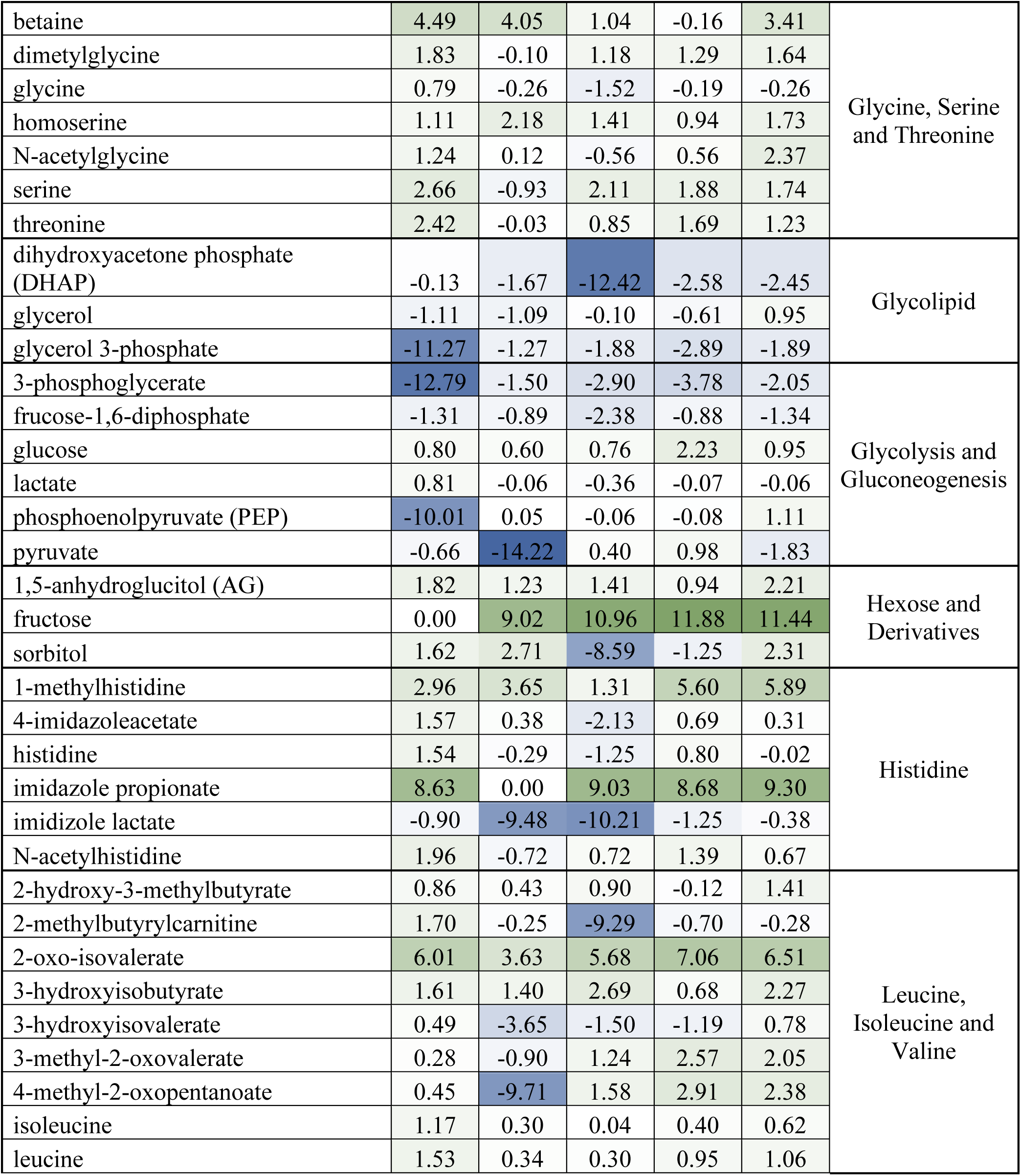
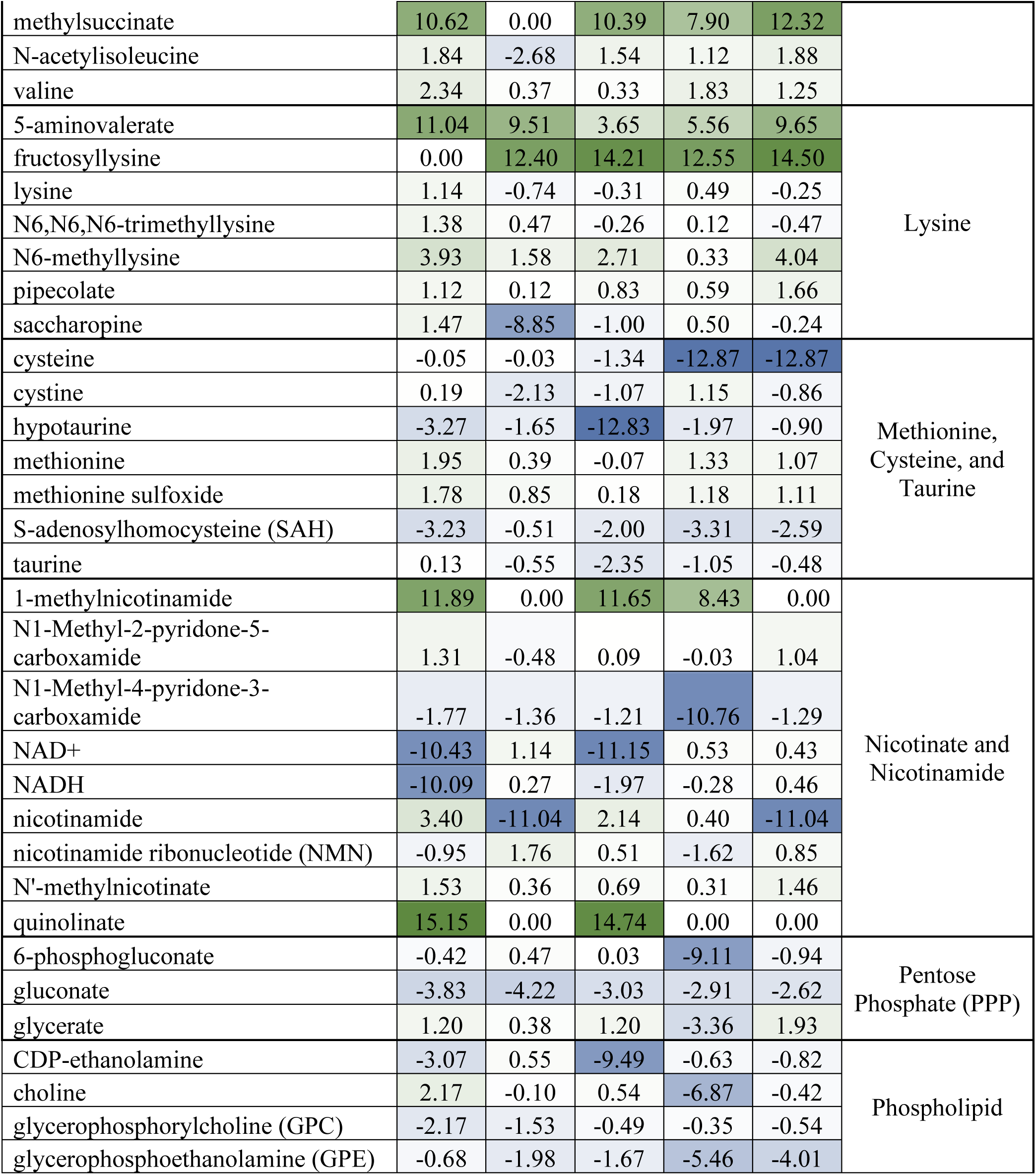
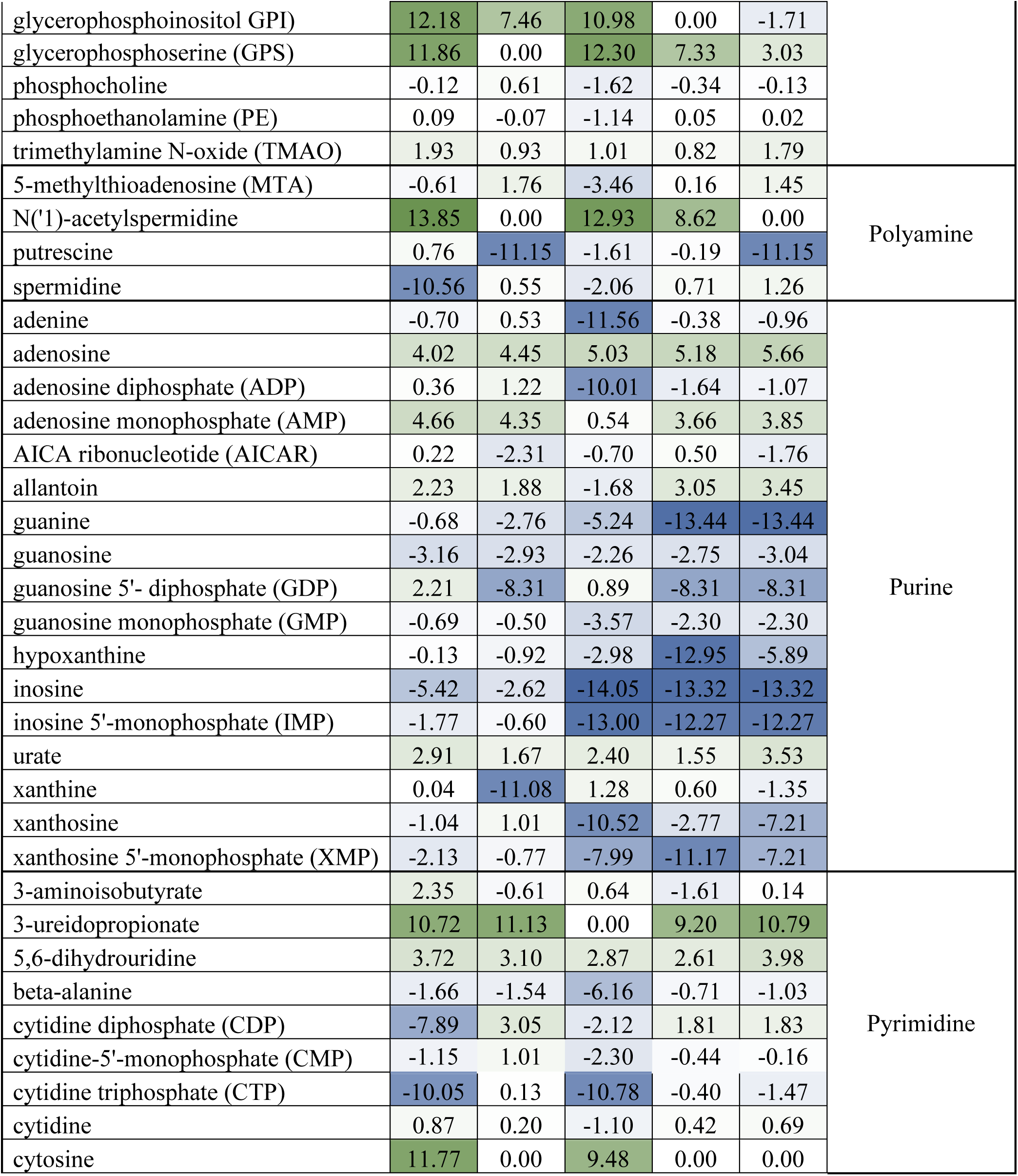
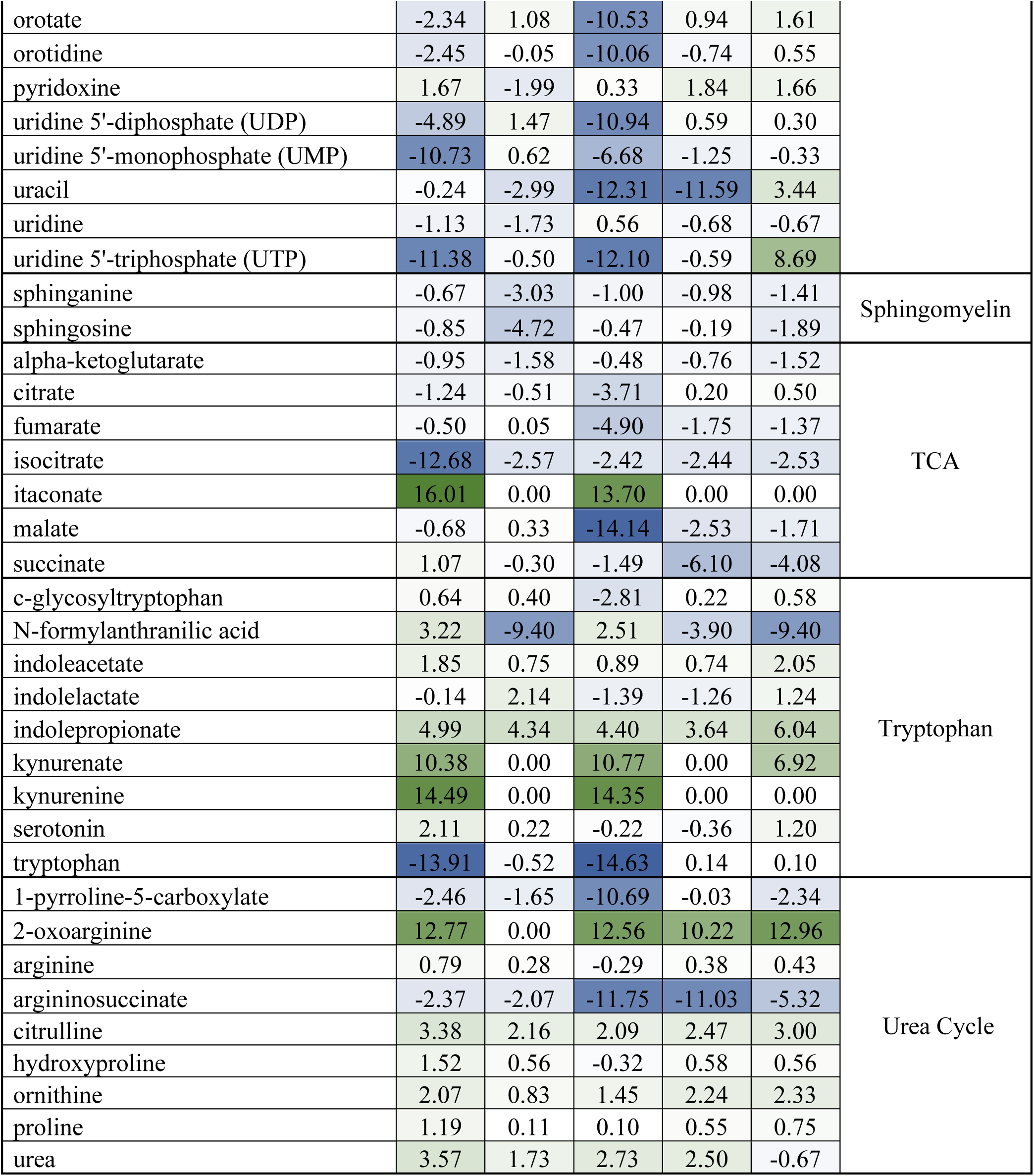
Relative Concentrations of Select Pathway Metabolites for Each Polarized Phenotype Compared to the Parent, Resting MΦ Phenotype (M0). Pathway categories and metabolites correspond to the data pictures in Figure 9. Heatmap scale (-8 to 8) is colored from blue to green.

